# HIPPO signaling competency links human stromal cell states to bone forming potential and osteogenic responsiveness

**DOI:** 10.64898/2026.11.14.766622

**Authors:** Atenisa Caci, Nicholas Ditzel, Sebastian Zanner, Moustapha Kassem, Alexander Rauch

## Abstract

Skeletal stromal populations contribute to bone maintenance and repair, but how their molecular heterogeneity relates to bone forming potential remains incompletely understood. Whether shared transcriptional programs link this functional diversity to osteogenic responsiveness remains unresolved. We combined single-cell RNA sequencing of four human stromal clones with contrasting bone forming capacities, functional assays, and primary human stromal datasets. High bone forming clones shared oxidative metabolism and signaling-associated programs despite clone-specific expression patterns. Metabolic flux measurements supported these differences, whereas mixed implants provided no evidence of paracrine transfer of bone forming potential. We therefore examined signaling-associated competency states and identified a HIPPO competency signature enriched in high bone forming clones despite lower canonical YAP/TAZ target-gene expression. Unlike target-gene expression, competency-associated expression remained comparatively stable following pathway perturbation. Functionally, high bone forming clones showed greater osteogenic responses to both pharmacological activation and inhibition of YAP/TAZ-dependent transcription. The signatures also showed partially distinct distributions in primary human bone marrow stromal populations, and lower competency scores in bone and stromal cells were associated with osteoporosis. These findings link HIPPO competency to bone forming capacity and osteogenic responsiveness, suggesting that shared signaling-associated states provide a basis for understanding functional differences among stromal progenitors beyond marker-defined identity.

## Introduction

Skeletal stem and progenitor cells supply osteoblasts for bone maintenance and repair, but comprise heterogeneous, partially overlapping populations whose relationships and functional properties remain incompletely resolved (Melis, Trompet et al. 2025). These populations occupy different skeletal compartments, and their contributions to osteogenesis depend on developmental stage and injury context. Several cellular sources can contribute to repair, indicating that osteogenic potential is not restricted to a single progenitor identity (Trompet, Melis et al. 2024). In human bone marrow stromal cells, conventional osteoblast markers and *in vitro* differentiation assays do not consistently predict ectopic bone formation (Larsen, Frederiksen et al. 2010, Janicki, Boeuf et al. 2011). Distinguishing molecular features associated with bone forming potential therefore requires linking cellular characterization to functional measurements. Single-cell studies have resolved cellular diversity in human bone marrow and the endosteal compartment (Bandyopadhyay, Duffy et al. 2024, Chai, Lundberg et al. 2026), while profiling of cultured primary stromal cells has identified an ITGA11-marked population associated with osteogenic potential (Jamil, Madsen et al. 2026). In the latter study, ITGA11-based isolation enriched cells with different osteogenic capacities, whereas *ITGA11* knockdown did not alter the associated transcriptional signature, distinguishing a population marker from the program it identifies.

Comparative studies reveal both shared and distinct properties across skeletal progenitor populations. Selected marrow and periosteal progenitors exhibit substantial transcriptional similarity (Deveza, Ortinau et al. 2018), whereas osteochondral and perivascular stem-cell populations differ in lineage potential (Ambrosi, Sinha et al. 2021). Marker expression also overlaps, with LEPR-lineage marrow stromal cells expressing *Cxcl12* and *Grem1* (Mo, Guo et al. 2022). Furthermore, periosteal progenitors can generate LEPR+ marrow stromal cells during fracture repair (Jeffery, Mann et al. 2022). These findings indicate that marker-defined identities do not map directly onto distinct functional properties. The common and population-specific functions of these progenitors remain incompletely resolved (Melis, Trompet et al. 2025). To what extent their osteogenic potential reflects shared transcriptional programs is therefore an important open question. One possibility is that such programs encode the capacity to respond to osteogenic cues. Stromal progenitors are pre-programmed by their tissue microenvironment, while their differentiation involves signaling pathways and transcriptional regulators acting in a coordinated sequence (Rauch and Mandrup 2021).

HIPPO signaling provides a framework for investigating this relationship. Its transcriptional effectors YAP and TAZ integrate inputs from cell adhesion, mechanical forces, growth factors, and cellular metabolism (Yu, Zhao et al. 2015). YAP/TAZ connect mechanical properties of the cellular environment to osteogenic differentiation and regulate the balance between osteogenic and adipogenic programs (Hong, Hwang et al. 2005, Dupont, Morsut et al. 2011). Beyond lineage choice, they support the expansion and differentiation of osteoblast precursors during fracture repair (Kegelman, Nijsure et al. 2021). Their effects are nevertheless stage-dependent, as ablation of YAP/TAZ in osteoblast progenitors enhanced differentiation while deletion in mature osteoblasts and osteocytes reduced bone formation (Xiong, Almeida et al. 2018). Together, these studies show that the osteogenic consequences of YAP/TAZ regulation depend on the cellular context. Whether stromal cells with different bone forming capacities also differ in their competency to respond to pathway modulation remains unresolved.

To address this question, we combined single-cell transcriptomics of high and low bone forming human stromal clones derived from a common parental population (Larsen, Frederiksen et al. 2010) with functional assays and comparisons to primary human stromal cell datasets. We identified shared signaling-associated programs in high bone forming cells and distinguished a HIPPO competency signature from canonical target-gene expression. This HIPPO competency signature was associated with bone forming potential, osteogenic responsiveness, and osteoporosis status in primary stromal cultures.

## Results

### Cultured hBMSCs exhibit distinct bone forming capacities in vivo

To assess functional heterogeneity within human bone marrow stromal cells (hBMSCs), we reanalyzed previously established clonal cell lines derived from hBMSC-TERT4 cells (Simonsen, Rosada et al. 2002, Larsen, Frederiksen et al. 2010) in an ectopic bone formation assay. Consistent with their previously reported phenotypes, histological analysis affirmed marked differences in the capacity to form bone, with human-specific vimentin staining identifying osteocytes of human origin within the newly formed bone (Fig. 1A). Based on the quantification of bone formed we distinguished high bone forming (HBF; e.g., DD8 and AD10) and low bone forming (LBF; e.g., CB4 and CD8) clones (Fig. 1B). Although all lines retained a broadly fibroblastic appearance in culture (Fig. 1C), quantitative analysis of nuclear morphology revealed smaller, more elongated nuclei in DD8 as a HBF clone compared to CB4 as LBF clone (Fig. 1D, E). The lower width-to-length ratio is consistent with previous findings linking elongated nuclear geometry to enhanced osteogenic differentiation and matrix mineralisation in primary human bone marrow stromal cultures, and to greater *in vivo* bone forming capacity in other lines derived from the same parental hBMSC-TERT4 cells (Kowal, Schmal et al. 2020). Given our previous finding that transcriptional features distinguish osteogenic potential beyond cell morphology in primary human stromal cells (Jamil, Madsen et al. 2026), we next used single-cell RNA sequencing to investigate molecular differences between and within the clonal lines.

**Figure 1:**
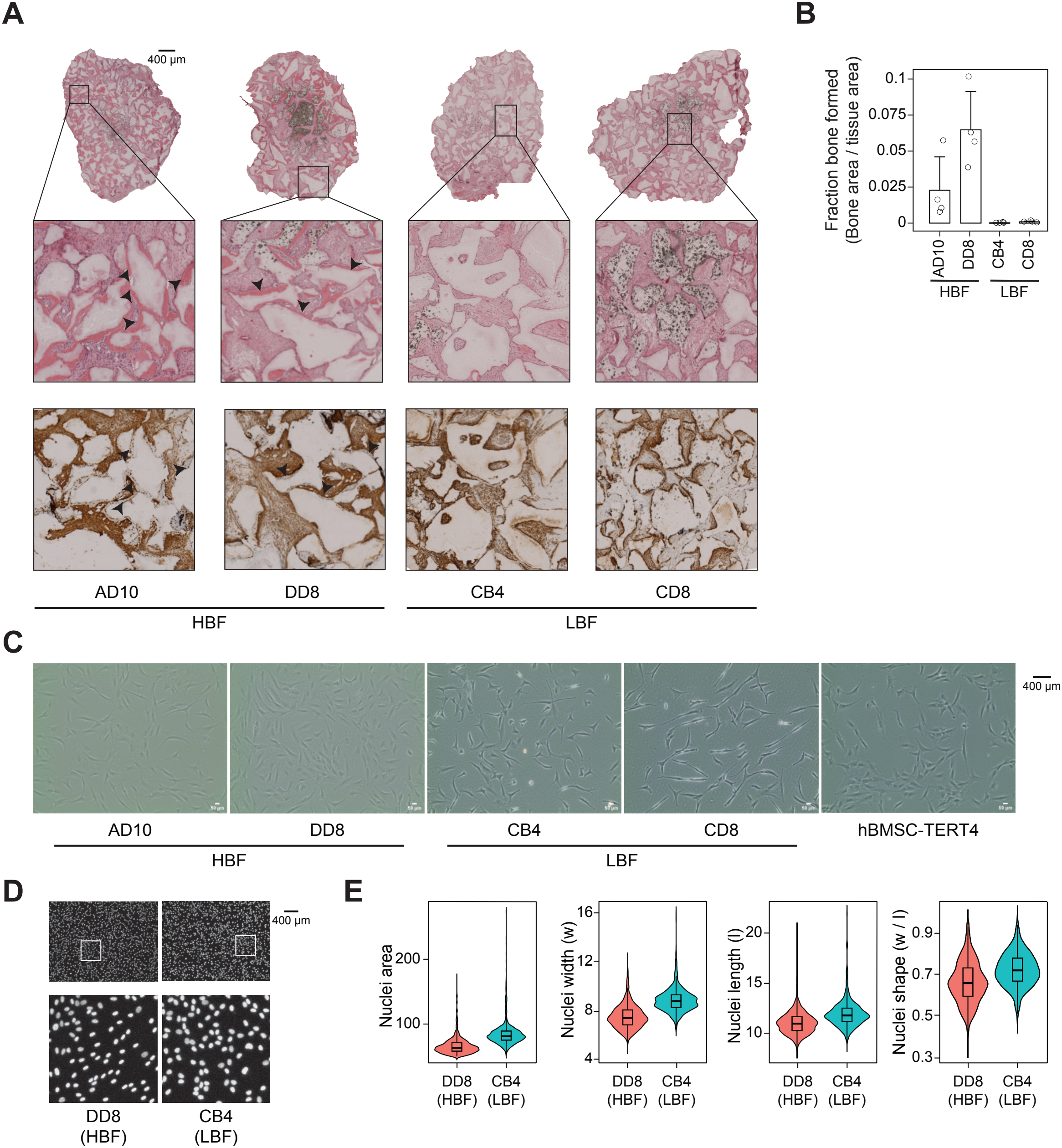
hBMSC-TERT4 clones with varying bone-forming potential. **A)** H&E staining (upper panel) and immune staining against human-specific vimentin (lower panel) of close-by sections from ectopic bone formation assays using hBMSC-TERT4 clones (AD10, DD8, CB4, CD8) in immune-deficient mice. Black arrowheads indicate areas of bone formation with deposition of vimentin-positive osteocytes. **B)** Bar plot quantifying bone as a percentage of bone area over tissue area from ectopic bone formation assays in 1A. LBF – Low bone forming, HBF – high bone forming. n = 4. **C)** Phase contrast microscopy at 10 x magnification of parental hBMSC-TERT4 cells and clonal lines with low and high bone forming capacity. **D)** DAPI stained nuclei from DD8 and CB4 cells at 4x magnification, with zoom on inlets. **E)** ImageJ-based quantification of nuclei from Fig. 1D. All parameters showed a p-value below 2.2 x 10^-16^ using a Wilcoxon rank-sum test. DD8: 3012 objects, CB4: 1467 objects.

### Intra-clonal transcriptional heterogeneity overlaps with clone-specific expression patterns

To investigate molecular differences between and within the clonal lines, we performed single-cell RNA sequencing of parental hBMSC-TERT4 cells and the four clonal cell lines. Joint UMAP embedding and clustering revealed 11 transcriptional states (Fig. 2A). Each clonal line contributed cells to multiple clusters, demonstrating intra-clonal heterogeneity, while AD10 and DD8 occupied broadly overlapping regions of the transcriptional landscape (Fig. 2B). Established stromal and skeletal progenitor markers suggested by the international society of cellular therapy (*THY1* encoding CD90, *NT5E* encoding CD73, *ENG* encoding CD105 (Dominici, Le Blanc et al. 2006)), from lineage tracing studies in mice (*LEPR (Zhou, Yue et al. 2014), CXCL12* (Omatsu, Sugiyama et al. 2010), PRRX1 (Liu, Li et al. 2022), *GLI1* (Shi, He et al. 2017), and *GREM1* (Worthley, Churchill et al. 2015)), from our previous study using cell surface marker expression to predict osteogenic capacity of primary stromal cells (*PDGFRA, NGFR* encoding CD271, *MCAM* encoding CD146, and *SDC2* encoding CD362 (Kowal, Moller et al. 2021)), as well as markers of more mature skeletal cells (*SP7* and *ADIPOQ* (Maes, Kobayashi et al. 2007, Zhong, Yao et al. 2020, Liao, Koehnken Sawall et al. 2025)) exhibited variable expression across clonal lines (Fig. 2C) and clusters (Fig. 2D). Here, the two HBF clones showed similar expression patterns as the parental cells, and *THY1* and *SDC2* clustered distinctly at the clonal and cluster level. Differential expression analysis identified cluster-enriched marker genes and clone-specific transcriptional signatures (Fig. 2E,2F). Six of the 11 clusters showed substantial overlap between cluster-enriched and clone-enriched genes, indicating that clonal identity contributes strongly to the signatures defining these clusters (Fig. 2F). Consistently, most cluster markers retained significant enrichment in only one or two cell lines (Fig. 2G). Nevertheless, selected markers showed conserved cluster enrichment across clones, such as the proliferation-associated cluster 4, as well as 3 and 9 (Fig. 2G, H). Separate clustering of each clone (Fig. 2I) further revealed intra-clonal subpopulations whose marker genes overlapped to varying degrees with those identified in the combined analysis (Fig. 2J). Thus, the transcriptional heterogeneity among the clonal cells is due to strong clone-specific patterns as well as conserved cluster-associated programs that can be recovered across the clonal lines.

**Figure 2:**
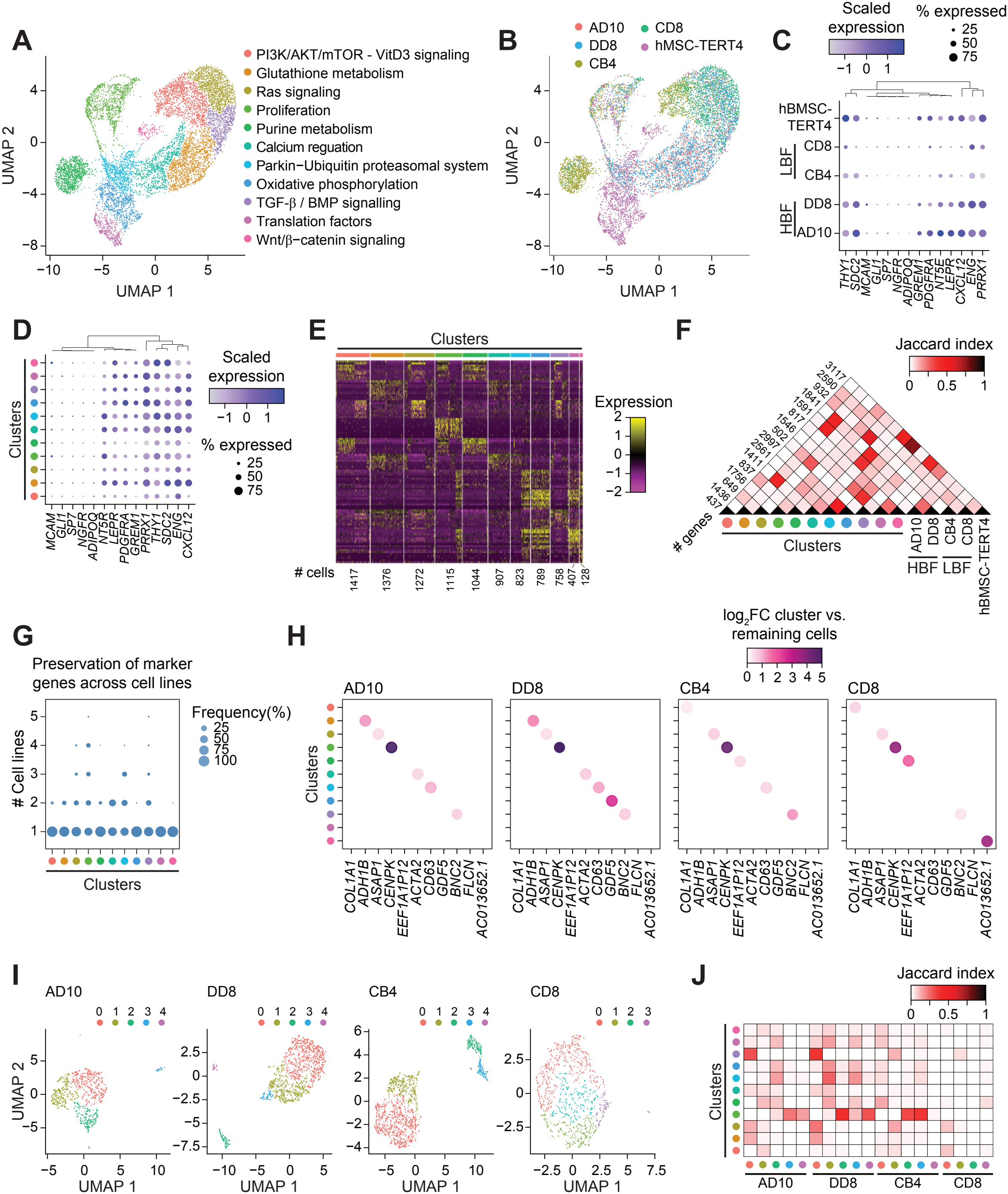
Intra-clonal transcriptional heterogeneity spans and overlaps inter-clonal variation in hBMSC-TERT4 cells. **A)** UMAP plot of 10036 cells derived from parental hBMSC-TERT4 cells and clonal lines with low and high bone forming capacity. Cluster annotations based on WikiPathway enrichment in Fig. 3B. **B)** UMAP-plot showing sample distribution from Fig. 2A. **C)** Dot plot of selected stromal cell progenitor markers across the clonal lines as in Fig. 2B. **D)** Dot plot of selected stromal cell progenitor markers across the identified clusters in Fig. 2A. **E)** Heat map showing scaled expression levels of the top 10 genes with enriched expression for the individual clusters. **F)** Heat map indicating Jaccard index for the pairwise overlap of cluster-specific and cell line-specific enriched marker genes. Enriched genes are defined as significantly (padj < 0.05) upregulated genes comparing the cells of interest against all remaining cells. The number of enriched cluster marker genes is indicated for each group. **G)** Dot plot showing the proportion of cluster-enriched genes that remain significantly enriched across individual samples. **H)** Color-coded dot plot showing the fold changes of globally defined enriched cluster marker genes across individual clonal lines. Fold changes were computed per line by comparing cells in the cluster of interest to all remaining cells within the same line. **I)** UMAP representations of scRNA-seq data from the 4 clones individually. **J)** Heat map indicating Jaccard index for the pairwise overlap of cluster-specific genes from the combined scRNA-seq object in Fig. 2A and the scRNA-seq objects of the individualized clones in Fig. 2I.

### High bone forming clones share cluster compositions associated with signaling and oxidative metabolism

To investigate whether transcriptional heterogeneity relates to bone forming potential, we compared the distribution of cells across clusters in the four clonal lines. The HBF clones AD10 and DD8 exhibited similar cluster compositions, whereas the LBF clones CB4 and CD8 differed from both the HBF clones and each other (Fig. 3A). Pathway enrichment analysis linked clusters preferentially represented in HBF clones to various growth factor signaling, oxidative phosphorylation, mitochondrial electron transport, and fatty acid metabolism. Clusters preferentially represented in LBF clones were associated with glycolysis, bone development, cell migration, and amino acid metabolism (Fig. 3B-D). Given the prominent metabolic signatures and expression of glycolysis and oxidative phosphorylation-associated genes, we assessed the metabolic profiles of DD8 (HBF) and CB4 (LBF) cells using Seahorse XF assays. Mitochondrial stress testing revealed significantly increased maximal respiration in DD8 cells compared to CB4 cells, while basal respiration showed only minor differences (Fig. 3E, F). In contrast, glycolytic stress testing demonstrated elevated basal and compensatory glycolysis in CB4 cells (Fig. 3G, H). Integration of OCR and ECAR measurements confirmed distinct metabolic states, with DD8 cells displaying a more oxidative phosphorylation phenotype and CB4 cells exhibiting enhanced glycolytic activity (Fig. 3I). Together, these findings identify similar cluster compositions in HBF clones and demonstrate metabolic differences between representative HBF and LBF lines consistent with their transcriptional profiles.

**Figure 3:**
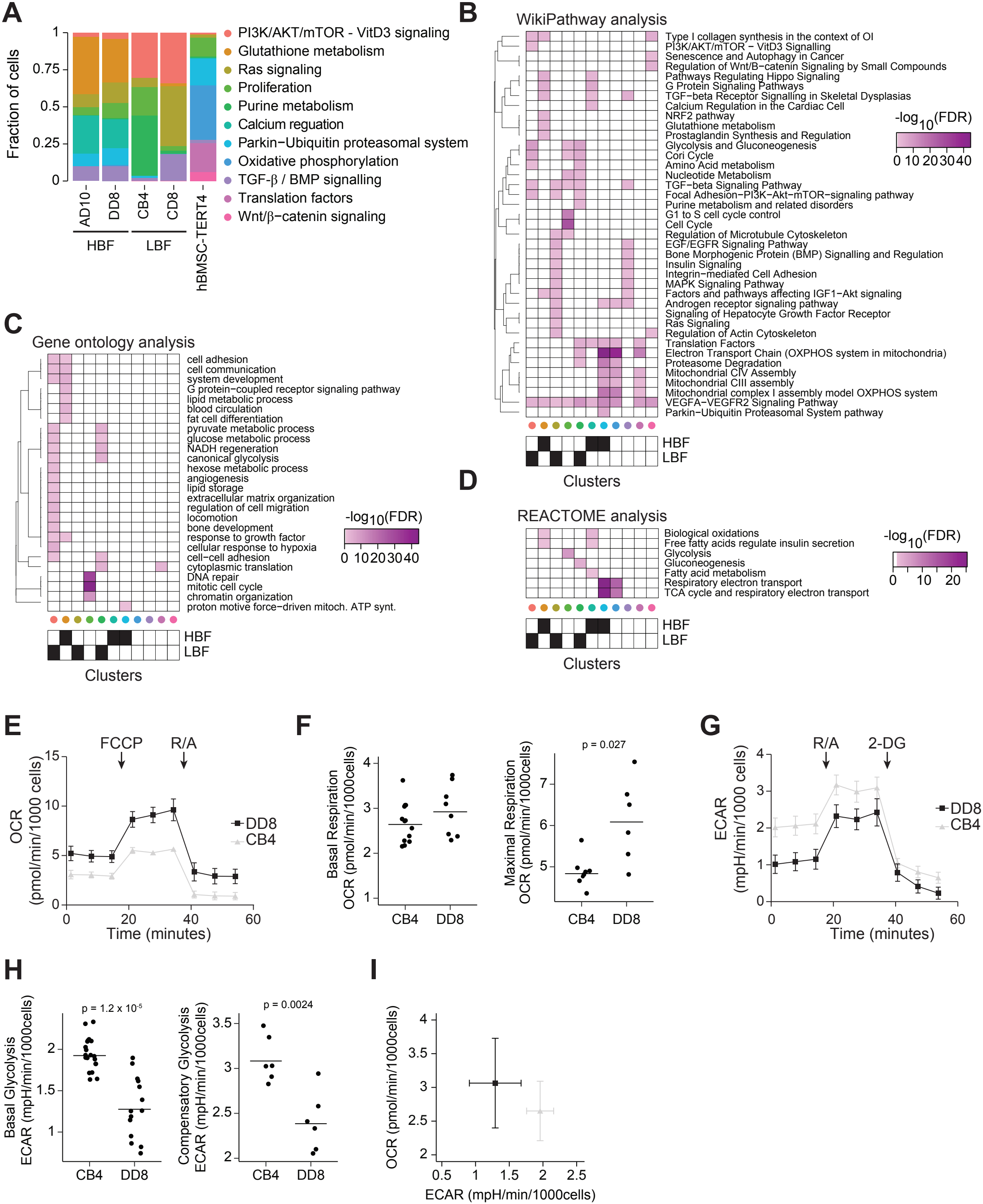
Subpopulation composition of hBMSC-TERT4 clonal lines aligns with clone-specific metabolic profiles. **A)** Stacked bar plot quantifying distribution of cells among clusters for each cell line. **B)** Wiki-pathway analysis for enriched cluster marker genes (cutoff Benjamin-Hochberg p-adj < 0.01). Low- and high-bone forming-selective clusters are indicated. **C)** Gene ontology analysis for enriched cluster markers (cutoff Benjamin-Hochberg p-adj < 0.01). Low- and high-bone forming-selective clusters are indicated. **D)** Reactome pathway analysis for enriched cluster markers (cutoff Benjamin-Hochberg p-adj < 0.01). Low- and high-bone forming-selective clusters are indicated. **E)** Oxygen consumption rate (OCR) measured by Seahorse assay in low (CB4) and high (DD8) bone forming cells during mitochondrial stress test (FCCP – R/A). Time course of OCR normalized to cell number, n > 8. **F)** Quantification of basal respiration (left, n > 8) and maximal respiration (right, n > 6) derived from Seahorse OCR measurements shown in 3E. Lines represent mean, unpaired Student’s T-test. **G)** Extracellular acidification rate (ECAR) measured by Seahorse assay in CB4 and DD8 cells during glycolysis stress test (R/A – 2DG). Time course of ECAR normalized to cell number, n > 12. **H)** Quantification of basal glycolysis (left, n > 12) and compensatory glycolysis (right, n > 6) derived from Seahorse ECAR measurements shown in 3G. Lines represent mean, unpaired Student’s T-test. **I)** Scatter plot of OCR versus ECAR summarizing metabolic phenotype of CB4 and DD8 cells. Mean values with error bars representing standard deviation, n > 6.

### Cluster-associated transcriptional programs relate to stromal differentiation and human bone biology

To evaluate the biological relevance of the identified cluster-specific gene signatures, we compared the cluster marker genes with independent datasets related to heterogeneity of primary stromal cells, differentiation and disease-associated expression profiles, as well as human genetics of bone mineral density. Our recently identified *ITGA11*-marked population (cluster 0) that demarcates *in vitro* mineralization and ectopic bone forming capacity of primary stromal cells (Jamil, Madsen et al. 2026) was enriched for marker genes of all HBF-associated clusters (Fig. 4A). Likewise, cluster 3 of the primary stromal cells, which shared marker genes with cluster 0 but was neither linked to osteogenic nor adipogenic potential, strongly overlapped with marker genes of all LBF-associated clusters. With focus on dynamic gene expression during differentiation of hBMSC-TERT4 cells (Rauch, Haakonsson et al. 2019), we found HBF clusters to be enriched for genes showing transcriptional activation during both osteoblast and adipocyte differentiation (Fig. 4B). Linking the cluster genes to human genetic traits of bone mass (Morris, Kemp et al. 2019), we found almost all cluster markers to show proximity-based enrichment for variants associated with estimated bone mineral density, with HBF clusters showing a quantitatively stronger association than LBF clusters (Fig. 4C). We next assessed the relationship of cluster markers to gene expression in osteoporotic subjects and bone forming capacity of primary cells. Microarray profiles from iliac crest biopsies of subjects with and without osteoporosis (Jemtland, Holden et al. 2011) suggested that HBF-associated cluster markers decline while LBF-associated ones increase expression in osteoporotic subjects (Fig. 4D). Comparison with bone marrow-derived stromal cells classified by the ability to form ectopic bone (Janicki, Boeuf et al. 2011) revealed a less uniform relationship; markers of one HBF-associated cluster were expressed at higher levels in bone forming cultures, while markers of the other two HBF-associated clusters as well as the LBF-associated clusters declined in bone-forming competent cells (Fig. 4E)

**Figure 4:**
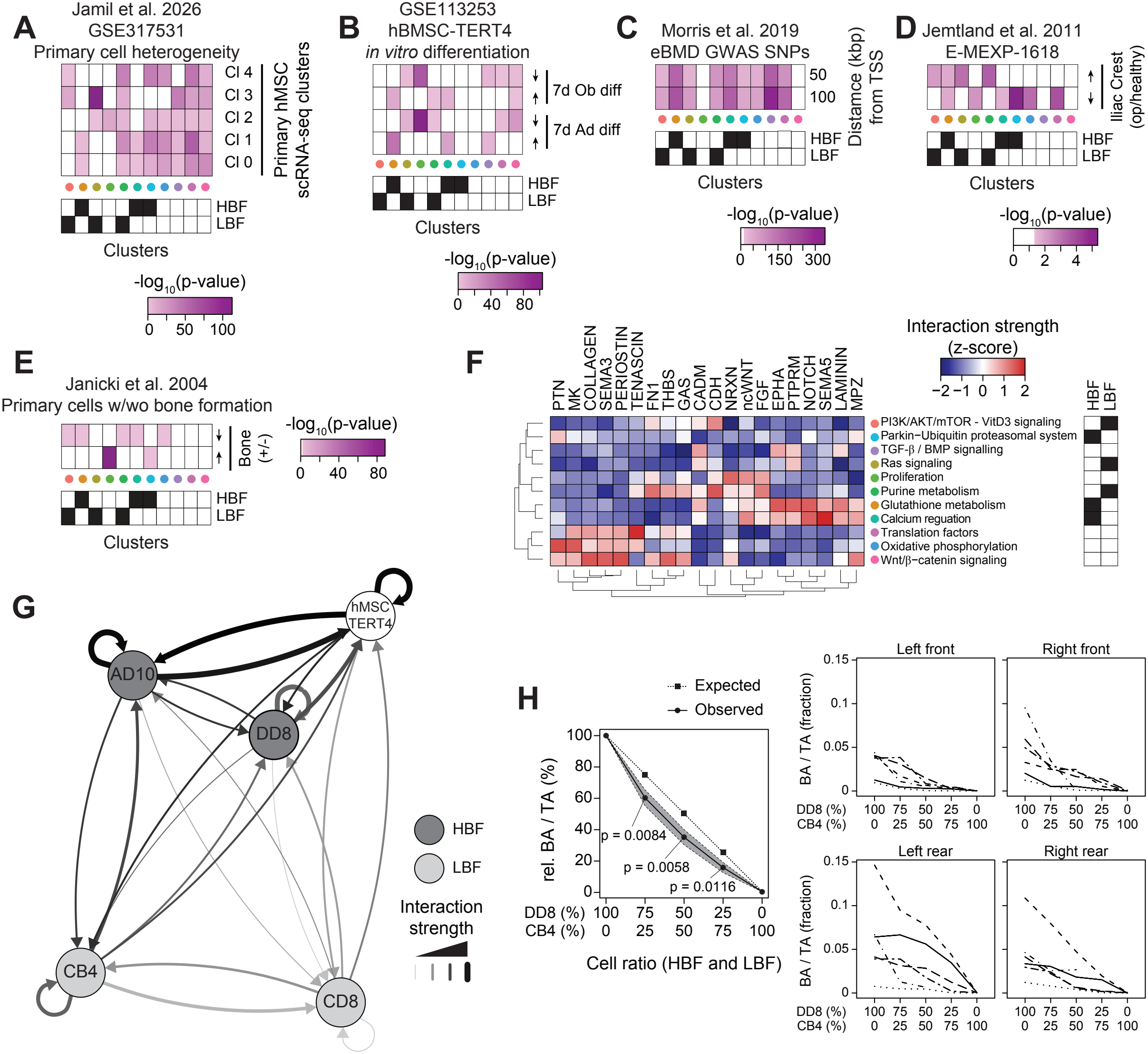
Cluster-enriched genes associate with bone biology and interaction profiles but fail to transfer bone forming potential in a paracrine setting. **A)** Hypergeometric-based enrichment (cutoff p-val < 0.05) of cluster-enriched marker genes for genes showing cluster-specific expression patterns in primary stromal cells (Jamil, Madsen et al. 2026). Here, Cl 0 was highly associated with *in vitro* osteoblast differentiation capacity. Low- and high-bone forming-selective clusters are indicated. **B)** Hypergeometric-based enrichment (cutoff p-val < 0.05) of cluster-enriched marker genes for genes showing differential expression (DESeq2-derived padj < 0.01) after 7 days of osteogenic or adipogenic differentiation in hBMSC-TERT4 cells (Rauch, Haakonsson et al. 2019). Low- and high-bone forming-selective clusters are indicated. **C)** Heat map showing the distance-dependent and chi-square-based enrichment of eBMD-associated SNPs (Morris, Kemp et al. 2019) near enriched cluster markers (cutoff p-val < 10^-5^). Low- and high-bone forming-selective clusters are indicated. **D)** Hypergeometric-based enrichment (cutoff p-val < 0.05) of cluster-enriched marker genes for genes showing differential expression (limma-derived p-value < 0.005) in iliac crest bone biopsies of subjects with osteoporosis compared to controls (Jemtland, Holden et al. 2011). Low- and high-bone forming-selective clusters are indicated. **E)** Hypergeometric-based enrichment (cutoff p-val < 0.05) of cluster-enriched marker genes for genes showing differential expression in primary cells with and without ectopic bone forming capacity (Janicki, Boeuf et al. 2011). Low and high bone forming-selective clusters are indicated. **F)** Heat map showing inferred cell–cell communication pathways based on CellChat analysis. Color scale represents interaction strength (z-score) for ligand–receptor signaling families across clusters. Low- and high-bone forming-selective clusters are indicated. **G)** Network representation of inferred cell–cell communication between hBMSC-TERT4 clonal lines based on CellChat analysis. Nodes represent individual cell lines and edges indicate predicted interactions, with edge thickness corresponding to interaction strength. Node color denotes high bone forming (HBF) and low bone forming (LBF) clones. **H)** Ectopic bone formation assay of implants consisting of various ratios of cells from high (DD8) and low (CB4) bone forming clones. Left panel: Relative bone formation across 4 implantation sites (left and right, front and rear) and 6 biological replicates (n > 20). Black line shows mean with grey arear representing the standard deviation. One-sample Student’s T-test comparing the observed distribution to the expected value. Right panel: Line plots quantifying bone as fraction of tissue area for each individual implantation site. Lines indicate various biological replicates, n > 5.

These observations highlight that HBF-associated subpopulations exhibit gene programs associated with osteoblast differentiation, bone forming competency, genetic traits determinant of human bone mass, and declined expression in osteoporotic bone.

### HBF clones do not mediate bone formation through paracrine interactions

Because several HBF-associated cluster programs were related to signaling pathways and cell-matrix interactions, we next inferred cell–cell communication networks using CellChat analysis (Jin, Plikus et al. 2025). Distinct signaling interactions were identified across clusters, including pathways related to extracellular matrix signaling, semaphorins, WNT, FGF, and NOTCH signaling (Fig. 4F). At the clonal level, HBF clones showed stronger predicted outgoing interactions, indicating greater expression-based potential to provide signals to other cells (Fig. 4G). We therefore questioned whether maintenance of HBF gene signatures was due to autocrine signaling events and whether HBF cells could induce bone formation in LBF cells in a paracrine manner. We therefore performed ectopic bone formation assays with varying ratios of DD8 and CB4 cells. Mixed DD8/CB4 implants formed significantly less bone than expected from a simple additive model based on the proportions of the two clones (Fig. 4H), providing no evidence that HBF cells transfer their superior bone forming capacity to LBF cells through paracrine interactions. The reduction below the expected levels could reflect inhibition of HBF-mediated bone formation by LBF cells. Alternatively, efficient bone formation may require a critical number or local density of bone-competent cells, such that reducing the HBF fraction disproportionately limits bone formation. In this scenario, despite present in numbers, LBF cells cannot compensate because bone forming competence depends on intrinsic properties of HBF cells, or because LBF cells cannot respond adequately to signals provided by HBF cells. These possibilities are not mutually exclusive and cannot be distinguished by the mixed-implant assay alone.

### HBF clones exhibit signalling signatures associated with bone formation

Although inhibitory interactions of LBF cells may contribute to the mixed-implant phenotype, we next focused on signaling-associated transcriptional programs enriched in HBF cells that may reflect competency states related to bone forming potential. Those HBF-associated programs included calcium regulation, G protein signaling, TGFb signaling, HIPPO signaling, and the Parkin-ubiquitin pathway (Fig. 3B). Most genes within these signatures were not annotated as ligands or receptors in CellChat, indicating that the signatures extend beyond the components represented in the inferred communication networks (Fig. 5A). We next assessed whether their cumulative expression scores distinguished cells from HBF and LBF clones. Calcium regulation, HIPPO, TGFβ, and G-protein signaling signatures discriminated AD10 and DD8 from CB4 and CD8 cells (Fig. 5B), whereas the Parkin-ubiquitin pathway had no predictive power. To prioritize these programs for further investigation, we examined their overlap with markers of stromal cell populations in human bone marrow (Bandyopadhyay, Duffy et al. 2024) and putative causal genes associated with eBMD (Morris, Kemp et al. 2019). The HIPPO signaling signature was preferentially enriched in osteogenic and progenitor-like stromal cells (Fig. 5C) and contained a disproportionately high number of putative causal eBMD genes (Fig. 5D). Interestingly, the constituents of the HIPPO signaling signature have been implicated in developmental regulation, signaling input, propagation, integration, and mechanotransduction all with preferentially high expression in HBF clones (Fig. 5E). Together, these findings prioritized the HIPPO signaling gene signature for investigating the relationship between signaling competency and bone forming potential.

**Figure 5:**
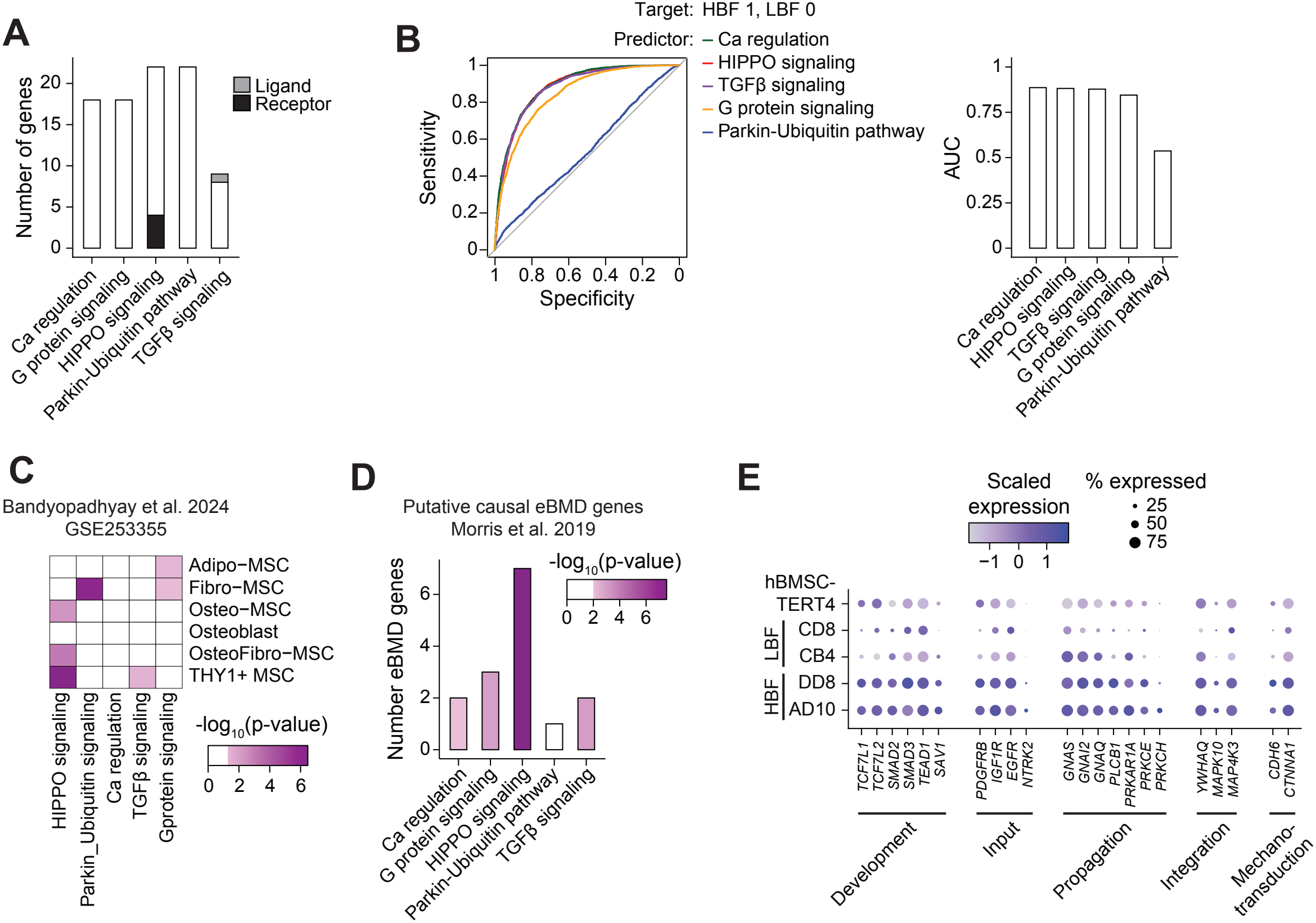
Signaling signatures demarcating high bone forming cells and implications in stromal cell identity and genetic traits of bone mass. **A)** Bar plot showing the number of genes in selected signaling pathways from Fig. 3B, indicating whether they are annotated as ligands or receptors in CellChat. **B)** ROC curve analysis predicting high and low bone forming cells using the scaled signature score of the indicated signaling pathways (left panel). Cells from AD10 and DD8 in the scRNA-seq object in Fig.2B were classified as 1, while CB4 and CD8 were classified as 0. Bar plot quantifying the area under the curve (right panel). **C)** Hypergeometric-based enrichment (cutoff p-val < 0.05) of signaling pathways in marker genes of defined stromal cell subpopulations from human bone (Bandyopadhyay, Duffy et al. 2024). **D)** Bar plot quantifying the number of putative eBMD genes among the signaling pathways, with color coding the hypergeometric-based enrichment. **E)** Dot plot highlighting the expression of a WikiPathway derived HIPPO signaling gene signature in hBMSC-TERT4 clones.

### HIPPO signaling competency distinguishes bone forming clones and stromal progenitor subsets

Since the HIPPO signaling signature does not comprise canonical HIPPO target genes such as *CCN1* (CYR61), *CCN2* (CTGF), or *ANKRD1* (Wang, Liu et al. 2017), we asked whether it reflects a signaling competency state distinct from downstream pathway output. To examine this distinction, we used published perturbation data across three human cell lines (GSE49384) and defined HIPPO target genes by the induction upon *NF2*/*LATS2* knockdown and subsequent normalisation upon *YAP1*/*WWTR1* knockdown (Fig. 6A). In contrast to HIPPO target genes, the HBF-associated HIPPO signature remained largely unchanged upon HIPPO pathway perturbation (Fig. 6A). We therefore termed the latter HIPPO competency signature, distinguishing its association with signaling capacity from the transcriptional output captured by the target signature.

**Figure 6:**
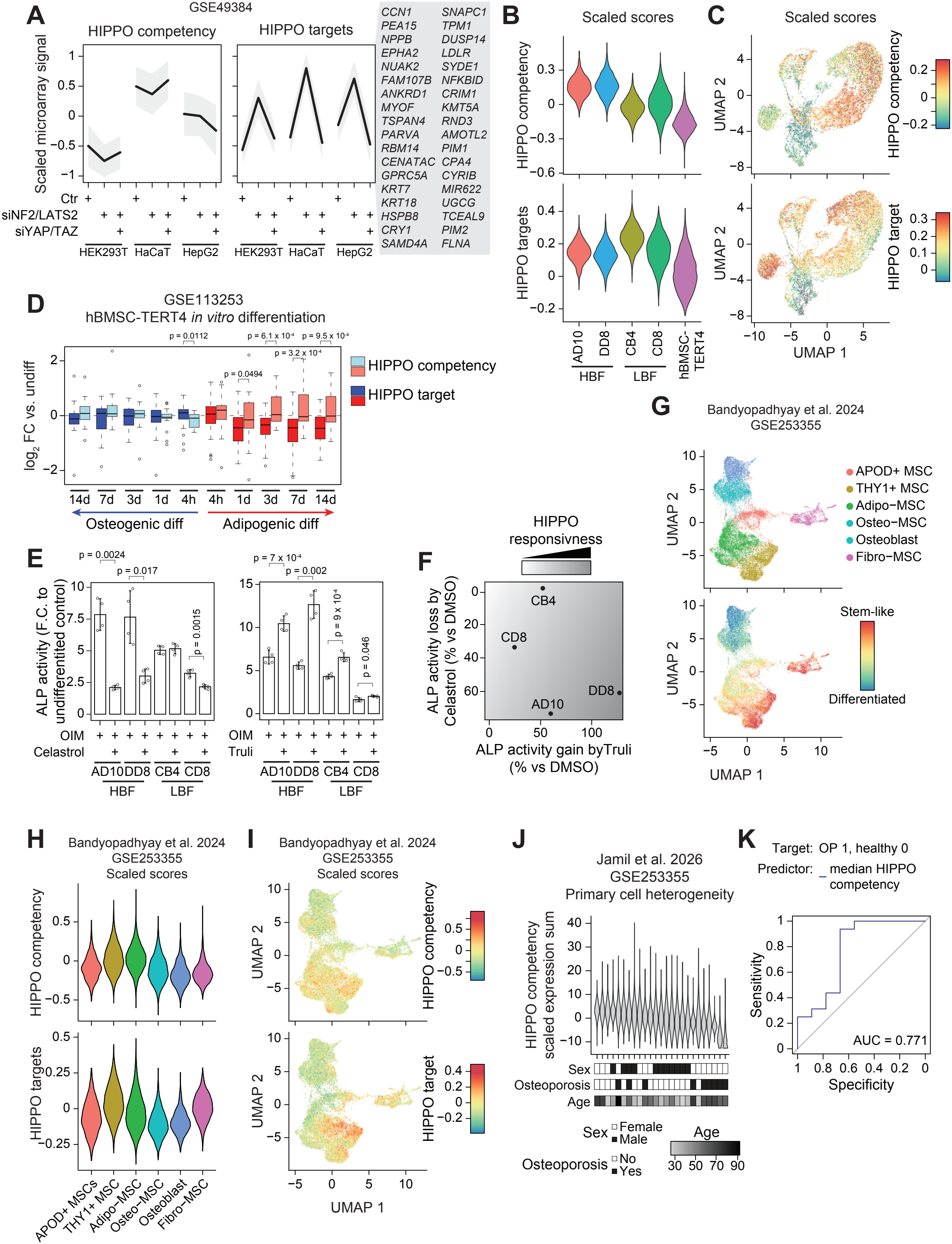
A HIPPO-signaling competency signature demarcates bone forming cells and distinct progenitors in human bone. **A)** Average scaled expression of the HIPPO competency (left panel) and a HIPPO target gene signature (right panel) under HIPPO signaling activation (*NF2*/*LATS2* knockdown) and inactivation (*NF2*/*LATS2* + *YAP1*/ *WWTR1* knockdown; *WWTR1* encoding TAZ) in various human cell lines. HIPPO target genes were defined by showing a cell line independent activation upon NF2/LATS2 knockdown and repression upon additional YAP/TAZ knockdown, genes are depicted to the right. Line showing the median and grey area the standard deviation. **B**) Summed scaled gene expression of the HIPPO competency (upper panel) and HIPPO target gene signature (lower panel) in the scRNA-seq data of hBMSC-TERT4 cells and the clonal lines. **C**) Feature plot highlighting cells with high and low scores of the HIPPO competency (upper panel) and HIPPO target gene signature (lower panel) in the scRNA-seq data of hBMSC-TERT4 cells and the clonal lines. **D**) Box plot quantifying changes in gene expression of the HIPPO competency and target gene signature in hBMSC-TERT4 cells upon osteogenic or adipogenic differentiation (Rauch, Haakonsson et al. 2019). Wilcoxon rank-sum test. **E**) Relative ALP activity (normalized to cell viability) of clonal cell lines at day 7 of osteogenic differentiation upon exposure to Celastrol (30 nM, left panel) or Truli (125 nM, right panel). ALP activity of each clonal line in the undifferentiated state was normalized to 1. Bars show the mean, and error bars show standard deviation. Unpaired Student’s T-test, n = 4. **F**) Scatter plot expressing the percentage of gained or lost ALP activity upon Truli or Celastrol exposure for the individual clones. Color shading represents strong susceptibility to altered ALP activity upon activation or repression of HIPPO signaling. **G**) UMAP plot (upper panel) and Feature plot (lower panel) of the stemness potential of stromal cells from human bone (Bandyopadhyay, Duffy et al. 2024). **H**) Summed scaled gene expression of the HIPPO competency (upper panel) and HIPPO target gene signature (lower panel) in the scRNA-seq data of stromal cells from human bone (Bandyopadhyay, Duffy et al. 2024). **I**) Feature plot highlighting cells with high and low scores of the HIPPO competency (upper panel) and HIPPO target gene signature (lower panel) in stromal cells of human bone (Bandyopadhyay, Duffy et al. 2024). **J**) Violin plot showing the sum of the scaled expression levels of the HIPPO competency signature across primary cells obtained from male and female donors with and without osteoporosis (Jamil, Madsen et al. 2026). Samples are arranged in declining order of the median. **K**) ROC-curve analysis for prediction of diagnosis of osteoporosis using the median expression of the HIPPO competency signature in Fig. 6J. Area under the curve (AUC) is indicated.

Projection of both signatures onto the clonal single-cell dataset revealed distinct expression patterns. HBF clones exhibited elevated HIPPO competency scores but lower target-gene scores than LBF clones (Fig. 6B, C), indicating that enrichment of the HIPPO competency signature was not necessarily linked to greater basal YAP/TAZ-dependent transcriptional output. Both HIPPO target and competency signature remain stable during osteogenic differentiation of hBMSC-TERT4 cells. In contrast, during adipogenic differentiation competency-associated genes increased expression while HIPPO target genes declined (Fig. 6D). Thus, the two HIPPO signatures capture distinct transcriptional features that can change independently during differentiation.

To test whether elevated expression of the HIPPO competency signature was associated with functional responsiveness, we assessed pharmacological pathway modulation during osteogenic differentiation. Celastrol and Truli, components repressing and activating YAP/TAZ dependent gene transcription (Nouri, Azad et al. 2019, Kastan, Gnedeva et al. 2021) were tested on the clones to modulate osteogenic differentiation *in vitro*. While Celastrol reduced and Truli activated ALP activity in osteogenic stimulated cells (Fig. 6E), HBF clones showed greater relative responses than LBF clones (Fig. 6F), supporting the notion that HIPPO competency reflects functional pathway responsiveness in terms of osteogenic differentiation.

To assess the *in vivo* relevance of these signatures, we mapped HIPPO competency and HIPPO target gene expression onto stromal cell populations from human bone marrow (Bandyopadhyay, Duffy et al. 2024). Both signatures displayed heterogeneous distributions across stromal subpopulations, with enrichment in distinct progenitor and osteogenic states (Fig. 6G, H). Interestingly, while both the HIPPO competency and HIPPO target gene signatures mapped predominantly to THY1+ MSC progenitors, their expression scores localized to partially distinct cellular subsets (Fig. 6I). Thus, both signatures show partially distinct expression patterns within the transcriptionally defined THY1+ MSC compartment, highlighting HIPPO pathway-related heterogeneity within this annotated population.

Finally, analysis of scRNA-seq data from cultured primary human stromal cells (Jamil, Madsen et al. 2026) demonstrated substantial inter-individual variation in the expression of the HIPPO competency signature (Fig. 6J). Stratification by donor characteristics indicated that osteoporosis diagnosis, but not donor age or sex, was associated with altered HIPPO competency states. Consistently, ROC analysis demonstrated moderate discrimination between osteoporotic and non-osteoporotic samples based on HIPPO competency expression (AUC = 0.771; Fig. 6K). Together, these findings identify a HIPPO competency signature as a feature of functionally distinct stromal cell states associated with bone forming potential and osteoporosis status.

## Discussion

Our study links differences in stromal bone forming capacity to distinct transcriptional programs whose biological relevance extends beyond the clonal model. Despite substantial heterogeneity within each clone, HBF clones showed more similar cluster compositions than LBF clones, suggesting greater consistency in the transcriptional organization associated with high bone forming capacity. HBF-associated programs overlap with an osteogenic primary stromal population, differentiation-induced genes, and genes reduced in bone and primary cells from osteoporotic subjects. In contrast, LBF-associated programs overlap with a different primary stromal population, differentiation-repressed genes, and genes elevated in osteoporotic bone. These contrasting associations support different biological properties of HBF and LBF cells, although their signatures do not uniformly distinguish bone forming capacity across independent data sets. Within this broader functional heterogeneity, we identify HIPPO signaling competency as a shared feature of HBF clones that is distinct from basal pathway output and associated with osteogenic responsiveness.

### Clonal stromal heterogeneity includes shared and clone-specific transcriptional programs

Stromal cell (or MSC) heterogeneity has become a central concept in skeletal biology, with recent single-cell and lineage tracing studies identifying multiple osteogenic, adipogenic, fibroblastic, and progenitor-like stromal populations in both mouse and human bone marrow. Lineage tracing studies in mice identified *Lepr*^*+*^ stromal cells as major skeletal stem and adipogenic progenitors within the adult marrow niche (Zhou, Yue et al. 2014), while additional studies described nestin-expressing progenitors (Mendez-Ferrer, Michurina et al. 2010), regeneration-activated *Cxcl12*-abundant reticular (CAR) cells (Matsushita, Nagata et al. 2020), periosteal *Acta2*^*+*^ (Matthews, Wee et al. 2020) and *Ctsk*^*+*^ cells (Debnath, Yallowitz et al. 2018), *Grem1*^*+*^ osteochondroreticular stem cells (Worthley, Churchill et al. 2015), growth and growth plate associated *Col2a1*^*+*^, *Pthrp*^*+*^, and *Gli1*^+^ skeletal stem and progenitor populations (Ono, Ono et al. 2014, Shi, He et al. 2017, Mizuhashi, Ono et al. 2018), and *Adipoq*^*+*^ marrow adipogenic lineage precursors (MALPs) (Zhong, Yao et al. 2020). In parallel, single-cell atlases of mouse bone further subdivided stromal populations into *Cxcl12*^*+*^, osteogenic, fibroblastic, and adipogenic subsets (Baryawno, Przybylski et al. 2019, Tikhonova, Dolgalev et al. 2019), while more recent human studies proposed hierarchical relationships between progenitor-like THY1+ MSCs and lineage-committed stromal states (Bandyopadhyay, Duffy et al. 2024). These approaches provide complementary descriptions of stromal diversity, although the relationships between marker-defined populations, transcriptional states, and functional potential remain incompletely resolved (Zhu, Ding et al. 2022, Weldon, Longaker et al. 2025).

Skeletal stromal populations identified through these approaches are often partially overlapping and context-dependent, making them difficult to reconcile into a single hierarchy. Their functional properties may reflect developmental origin, anatomical localization, and environmental context, alongside their current transcriptional state (Mendez-Ferrer, Bonnet et al. 2020, Li, Xu et al. 2022, Weldon, Longaker et al. 2025). Our clonal analysis addresses a related question within a common parental cell model: to what extent are transcriptional programs shared across individual clones, and how does this relate to their bone forming potential?

The HBF clones showed similar overall cluster compositions, whereas the LBF clones differed more strongly from each other despite sharing some cluster-associated features. However, similarity in cluster composition was distinct from conservation of cluster-defining genes: many markers identified in the combined analysis did not retain cluster enrichment across individual clones. Conversely, selected programs, particularly the proliferation-associated program, showed greater cross-clonal consistency. Thus, bone forming capacity may reflect shared transcriptional signatures superimposed on clone-specific expression programs. More broadly, the osteogenic competency of various marker-defined progenitor populations might likewise involve shared transcriptional programs, while differences in lineage markers reflect developmental origin, anatomical localization, and niche context.

This distinction is relevant to the interpretation of combined single-cell analyses, as sample identity, donor background, and integration strategy can influence inferred cellular states. Recent benchmarking studies have shown that batch effects, sequencing depth, sparsity, and sample heterogeneity can substantially affect single-cell differential expression analysis (Nguyen, Baik et al. 2023), while joint-clustering approaches emphasize the need to preserve subtle biological differences across samples rather than remove them as technical variation (Moller and Madsen 2023). Our findings suggest that clone-specific expression patterns and shared functional programs can contribute simultaneously to transcriptional clustering. This interpretation complements previous descriptions of skeletal stromal populations as partially overlapping and context-dependent rather than readily organized into a single hierarchy (Li, Xu et al. 2022, Zhu, Ding et al. 2022, Weldon, Longaker et al. 2025). Identifying shared signatures of osteogenic competency across these populations may therefore help connect their molecular diversity to a common functional capacity.

### Bone forming potential associates with distinct metabolic and signaling programs

The shared transcriptional features of HBF clones included programs associated with growth factor signaling and oxidative metabolism, whereas LBF-associated clusters showed enrichment for glycolytic programs. Functional measurements in DD8 and CB4 clones supported this metabolic distinction, revealing greater maximal mitochondrial respiration in DD8 and greater basal and compensatory glycolysis in CB4. These findings are consistent with the increase in mitochondrial oxidative metabolism reported during osteogenic differentiation (Chen, Shih et al. 2008). However, our measurements were obtained in undifferentiated cultures, suggesting that metabolic differences can precede osteogenic induction and accompany differences in bone forming potential. Since bioenergetic analysis was limited to one HBF-LBF pair, we interpret metabolism as a component of these differences, without establishing whether it contributes causally.

The relationships of these programs to independent data sets suggest that HBF and LBF cells differ in biological properties extending beyond metabolism. HBF but not LBF-associated signatures overlapped with an ITGA11-marked primary stromal population linked to mineralization and ectopic bone formation (Jamil, Madsen et al. 2026). Enrichment of HBF-associated genes among those induced during both osteogenic and adipogenic differentiation (Rauch, Haakonsson et al. 2019) further suggests that these programs reflect aspects of differentiation competency rather than osteoblast commitment alone. Their contrasting relationships to osteoporosis, with HBF-associated genes being reduced while LBF-associated genes being increased, also connect the clonal phenotypes to disease-associated transcriptional changes (Jemtland, Holden et al. 2011). However, the less consistent separation in primary cultures classified by ectopic formation (Janicki, Boeuf et al. 2011) shows that not all HBF-associated programs are shared across experimental models of bone forming capacity.

Beyond their metabolic differences, HBF clones shared transcriptional programs associated with several signaling pathways, and CellChat analysis suggested HBF cells have a greater potential to send signals to other cells, raising the possibility that they could promote bone formation in LBF cells. However, mixed implants formed less bone than expected from the proportions of the two clones. This finding is compatible with inhibitory effects of LBF cells or their lack of responsiveness, but also with a requirement for a critical number or local density of bone-competent cells. The assay therefore does not resolve whether limited bone formation reflects inhibitory interactions, insufficient cooperation among bone forming competent cells, or an inability of LBF cells to respond adequately to available signals. These possibilities motivated us to examine whether HBF-associated signaling programs reflect cellular competency states related to bone forming potential.

### HIPPO signaling competency links stromal transcriptional states to osteogenic responsiveness

HIPPO signaling integrates biochemical and mechanical cues that regulate cellular growth and differentiation (Yu, Zhao et al. 2015, Panciera, Azzolin et al. 2017). Mouse studies have established pro-osteogenic roles for YAP/TAZ in skeletal progenitor expansion, bone development, and fracture repair (Dupont, Morsut et al. 2011, Kegelman, Nijsure et al. 2021, Collins, Lang et al. 2024, Nijsure, Tobin et al. 2025). Human cell studies likewise demonstrate a supportive role in osteogenesis: YAP/TAZ mediate mechanically regulated differentiation of human stromal progenitors (Dupont, Morsut et al. 2011), restoration of YAP1 or TAZ rescues impaired osteogenesis following kindlin-2 depletion in placenta-derived stromal cells (Guo, Cai et al. 2018), and TAZ manipulation alters mineralization and bone forming outcomes in human adipose-derived stromal cells (Zhu, Wu et al. 2018). Our findings therefore extend previous demonstrations that YAP1/TAZ regulate human osteogenesis by linking HIPPO competency to differences in bone forming capacity and the magnitude of osteogenic responses to pathway modulation.

In developmental biology, competence describes the context-dependent ability of a cell to respond to inductive signals. Studies of regulatory landscapes and lineage priming illustrate how pre-existing cellular states can influence subsequent differentiation outcomes (Stergachis, Neph et al. 2013, Buenrostro, Corces et al. 2018, Meuleman, Muratov et al. 2020, Weinreb, Rodriguez-Fraticelli et al. 2020). Our findings suggest that a related principle may contribute to stromal osteogenic potential. The competency signature includes genes associated with signaling input, propagation, integration, and mechanotransduction, and its expression remained comparatively unchanged in data sets following perturbations that consistently altered YAP1/TAZ target-gene expression. Moreover, competency-associated expression increased during adipogenesis while target-gene expression declined, indicating that the signatures capture distinct aspects of cellular state. This increase also shows that the competency signature is not exclusive to osteogenic differentiation. Its association with bone forming potential may therefore reflect a broader signaling-related state whose functional consequences depend on the differentiation context.

The greater sensitivity of HBF clones to both Celastrol and Truli provides functional support for an association between HIPPO competency and the extent of osteogenic responsiveness. However, these experiments measure an osteogenic outcome rather than signaling activity directly, and pharmacological responses alone do not establish that the signature genes determine responsiveness. Whether individual signature components contribute causally to osteogenic responsiveness and bone forming capacity remains to be established.

### HIPPO competency in primary stromal cells and osteoporosis

The distinction between HIPPO competency and target-gene expression extends beyond the clonal model, with partially distinct expression patterns also observed within the THY1+ MSC compartment of human bone marrow *in vivo* (Bandyopadhyay, Duffy et al. 2024). Functional diversity within marker-defined stromal populations is supported by studies identifying osteogenic and mechanically responsive subsets among LEPR+ cells (Shen, Tasdogan et al. 2021), and reduced osteogenic regeneration following repeated activation of the LEPR-marked stromal lineage (Wu, Matsumoto et al. 2026). Signaling competency states may therefore provide an additional dimension for interpreting functional potential within progenitor populations and help explain differences that are not resolved by shared marker expression alone.

The association between lower HIPPO competency scores and osteoporosis in cultured primary stromal cells suggests that this transcriptional state may also be relevant to skeletal disease. Previous studies have linked skeletal aging and bone loss to impaired progenitor function, altered mechanosensation, and reduced regenerative capacity (Yang, Cao et al. 2023, Ambrosi, Taheri et al. 2025, Liu, Ye et al. 2026). Our findings raise the possibility that reduced signaling competency contributes to these functional changes, although the association does not establish whether lower signature expression reflects altered states within cells or differences in stromal subpopulation abundance. The observed donor-level discrimination is therefore evidence of a disease association rather than validation of a diagnostic marker. Testing whether HIPPO competency predicts osteogenic responsiveness in primary cells will be important for establishing its functional relevance beyond the clonal model.

### Limitations and future perspective

The four clonal lines originate from a single telomerase-immortalized parental population (Larsen, Frederiksen et al. 2010), limiting the extent to which their properties can be generalized to primary stromal cells across donors. Although comparisons with primary-cell datasets support the relevance of the identified programs, their functional relationship to HIPPO responsiveness remains to be tested in primary cells. The persistence of previously reported bone forming differences is consistent with maintenance of osteogenic potential during propagation, but does not establish how this potential is transmitted through cell divisions. Genetic differences, heritable epigenetic states, and self-reinforcing regulatory circuits remain possible explanations for the clonal phenotypes.

Our experiments associate HIPPO competency with osteogenic responsiveness but do not establish how this transcriptional state is generated or maintained. The pharmacological treatments probe the functional consequences of pre-existing competency, while the conditions required to establish that competency remain unresolved. Future studies should investigate whether defined biochemical or mechanical conditions can induce the coordinated transcriptional program and thereby confer greater responsiveness to subsequent pathway modulation. Testing whether acquisition of this state also enhances mineralization and bone formation would help establish its functional significance. A central question is whether modifying HIPPO competency can bring marker-defined stromal populations toward comparable osteogenic capacities. This would test our hypothesis that shared functional programs contribute to bone forming potential across populations distinguished by developmental origin, anatomical location, or marker expression. Determining whether an induced change persists after withdrawal of the intervention and subsequent cell divisions would further distinguish a transient enhancement of differentiation from a lasting change in osteogenic potential. Such studies could establish whether HIPPO competency is a modifiable property that helps explain—and potentially reduce—functional differences among stromal progenitor populations.

## Materials and methods

### Cell culturing

hBMSC-TERT4 cells (Simonsen, Rosada et al. 2002) and previously derived single-cell clones (Larsen, Frederiksen et al. 2010) were seeded in tissue culture flasks and expanded in culture medium consisting of minimum essential medium (MEM, 32561-094, Invitrogen) supplemented with 10% fetal bovine serum (FBS, 16000044, Life Technologies) and 1% penicillin/streptomycin (P/S, 15140-130, Invitrogen) and incubated at 37 °C with 5% CO_2_ and 95% humidity. hBMS-TERT4 cells were expanded at a split-ratio of 1:4 twice a week by trypsinization using 0.05% Trypsin-EDTA (25300062, Thermo Fisher Scientific) when reaching 80-90% confluency. Contamination with Mycoplasma was checked in the research unit every second month using a PCR test (N-GARDE Mycoplasma PCR Reagent set, BioNordika) on conditioned media, being at least 48 hours on the cells. All cultures reported here were negative for Mycoplasma.

Cells were treated with 2 different inhibitors, Celastrol (C0869-10MG, Sigma-Aldrich) or TRULI (SML3634-5MG, Merck). Inhibitors were dissolved in DMSO (187376, ILS) to obtain a stock solution of 5 mM, diluted in culture media, and refreshed upon media change. As control we used vehicle (DMSO) or untreated cells.

### In vitro cell differentiation

#### Osteoblast differentiation

hBMSC-TERT4 cells were seeded with a density of 40,000 cells/cm^2^ to achieve approximately 80% confluency the following day and expanded for 72 hours in culture medium. After 72 hours, culture medium was replaced with osteoblast induction medium consisting of MEM supplemented with 10% FBS, 1% P/S, 10 mM ß-glycerophosphate (50020-100, Sigma-Aldrich), 10nM dexamethasone (D4902, Sigma-Aldrich), 50µg/mL 2-phosphate ascorbic acid (A4544, Sigma-Aldrich) and 10nM 1,25-vitamin D_3_ (D1530, Sigma-Aldrich). Media was refreshed every 2-3 days.

#### Adipocyte differentiation

hBMSC-TERT4 cells were seeded with a density of 40 000 cells/cm^2^ to achieve approximately 100% confluency the following day and expanded for 72 hours in culture medium. After 72 hours, culture medium was replaced with adipocytic induction medium consisting of Dulbecco’s modified Eagle’s medium (DMEM, 41965062, Thermo Fisher Scientific) supplemented with 10% FBS, 1% P/S, 100nM dexamethasone, 3µg/mL insulin (I9278, Sigma-Aldrich), 1µM rosiglitazone (BRL, 71740, Cayman Chemicals) and 450µM 3-isobutyl-1-methylxanthine (IBMX, I5879-5G, Sigma-Aldrich). Media was refreshed every 2-3 days.

### Analysis of nuclear morphology

Cells were seeded at 20,000 cells/cm^2^ and cultured until reaching 80% confluence. Nuclei were stained with DAPI (62248, Thermo Fisher Scientific) for 10 minutes at room temperature while gently rotating and then washed 3 times for 5 minutes with PBS^-/-^. Images were captured by the Cytation imaging system at 4 × magnification. Images were analyzed utilizing ImageJ. Images were calibrated to 0.32 µm/pixel, and nuclei were segmented via Otsu thresholding in conjunction with watershed separation. Nuclear parameters such as area, perimeter, Feret’s diameter, ellipse-fitting metrics, and shape descriptors were assessed utilizing Analyze Particles. Objects with an area of ≥25 µm^2^ and a circularity ranging from 0.20 to 1.00 were included, whereas those in contact with the image edges were excluded.

Alkaline phosphatase (ALP) activity assay for measuring osteogenic differentiation Prior to measuring ALP activity, cell viability was measured through CellTiter-Blue assay according to the manufacturer’s protocol (Walzl, Unger et al. 2014) by replacing culture media with cell viability media (CellTiter-Blue (208657 (G8081), ILS Danmark) and culture medium, 1:5, respectively) and incubated at 37°C for 1 hour. Fluorescence intensity (579Ex/584Em) was measured with a FLUOstar Omega plate reader. For ALP activity, cells were washed twice with tris-buffered saline (TBS, (Lab44416.5000, Bie & Berntsen), pH=7,5), fixed in 3.7% formaldehyde (F1635, Sigma-Aldrich) – 90% ethanol for 30s and incubated with 1 mg/mL P-nitrophenylphosphate (dissolved in 50 mM NaHCO_3_ and 1 mM MgCl_2_ at pH=9.6, 71768-5G, Sigma-Aldrich) for 20 min. at 37°C. Absorbance was measured at 405 nm, and ALP activity was normalized to cell viability.

### Oil Red’O staining for measuring adipocyte differentiation

Lipid accumulation was measured through Oil Red’O staining. Differentiated cells were washed with PBS, fixed with 4% paraformaldehyde (PFA, D383004, Hounisen) in PBS at RT for 10 min, and washed with PBS. Fixed cells were stained with Oil Red’O solution (3:2 of 3mg/ml Oil Red’O (O0625, Sigma-Aldrich) in 100% isopropanol (I9516, Sigma-Aldrich) with H_2_O) for 1 hour at RT and rinsed with H_2_O. Images of lipid droplets were captured with a magnification of 10x using an Olympus IX50 microscope. Oil Red’O staining was extracted for quantification by adding isopropanol to the cells and incubated for 10 min at RT with agitation. Absorbance was measured at 500 nm.

### Ectopic bone formation

Assessment of bone forming capacity using an *in vivo* transplantation assay has been previously described in detail (Abdallah, Ditzel et al. 2008). Shortly, we used 500,000 hBMSC-TERT4 cells, first incubating with hydroxyapatite for 24 hours in syringes with growth media. We made 4 implants per mouse and compared bone formation to the same implantation sites in a paired manner. Implants were harvested after 8 weeks and fixed in 4% PFA for 24 hours, followed by decalcification in 0.5 M sodium formate for 72 hours. Implants were paraffine-embedded and sectioned at three different depths (100 µm apart). H&E-stained sections were quantified for bone using ImageJ by marking areas of bone normalized to the total area of the implant, and the presence of human cells in the xenograft was verified using immunohistochemistry with a human-specific vimentin antibody (abcam ab16700). As a deviation from the published protocol, NOD.Cg-*Prkdc*^scid^ *Il2rg*^tm1Sug^/JicTac mice have been used instead of NOD/LtSz-*Prkdc*^*scid*^ mice.

### Single cell RNA-sequencing and expression analysis

For single cell RNA-seq, hBMSC-TERT4 cells were trypsinized, spun down at 200 g, and washed twice in culture media. Cells were separated using a 40 µm mesh, washed 3 times in PBS with 0.04% non-acetylated BSA, and cell numbers were adjusted to 0.7 x 10^6^ cells per ml followed by a final transfer through a 40 µm mesh. Using 10^4^ cells as input material single-cell RNA-sequencing libraries were prepared according to the manufacturer’s instructions (10x Genomics). Cell gene count matrices were generated from raw sequencing data using the zUMI pipeline (Parekh, Ziegenhain et al. 2018) with human genome GRCh38 which yielded the following cell numbers: 2597 parental hBMSC-TERT4 cells; 6654 for AD10; 7257 for DD8; 10908 for CB4; and 42698 for CD8. Single-cell data were merged and filtered for high quality droplets (mitochondrial percentage less than 10%, gene count higher than 500) using the Seurat package (Butler, Hoffman et al. 2018) yielding a gene count matrix for 10036 high-quality cells. Clustering and quantification of gene signatures as summed scaled expression values were performed in Seurat (Butler, Hoffman et al. 2018).

Previously published data sets were downloaded from GEO: To define genes responsive to HIPPO signaling using GSE49384. To test genes for association with osteoporosis in elderly humans E-MEXP-1618 (Jemtland, Holden et al. 2011). To test differential expression during hBMSC-TERT4 cell differentiation into osteoblasts and adipocytes using GSE113253 (Rauch, Haakonsson et al. 2019). To explore defined stromal cell subpopulations of cultured primary cells using GSE317531 (Jamil, Madsen et al. 2026) as well as of human bone *in vivo* using GSE253355 (Bandyopadhyay, Duffy et al. 2024). Microarray data for GSE49384 and E-MEXP-1618 were processed with R packages GEOquery (Davis and Meltzer 2007), affy (Gautier, Cope et al. 2004), and limma (Ritchie, Phipson et al. 2015). For GSE113253, GSE317531, and GSE253355 processed count matrix and Seurat objects were used.

### Enrichment and cell-cell communication analysis

To infer biological functions of cellular subpopulations, we implemented open-source platforms WikiPathways (Pico, Kelder et al. 2008), Gene Ontology (Ashburner, Ball et al. 2000), Reactome (Ragueneau, Gong et al. 2026), and SPEED2 (Rydenfelt, Klinger et al. 2020) through GOseq (Young, Wakefield et al. 2010) or a hypergeometric test. Summary statistics from estimated bone mineral density GWAS (Morris, Kemp et al. 2019) were obtained from GEFOs (http://www.gefos.org/?q=content/data-release-2018). Single nucleotide polymorphisms (SNPs) were assigned to genes based on transcription start site annotations using GenomicRanges (Lawrence, Huber et al. 2013) and the *TxDb*.*Hsapiens*.*UCSC*.*hg19*.*knownGene* package applying variable genomic windows ranging from 50 to 2,500 kb. Enrichment analysis was performed using a Pearson’s chi-square test, comparing the proportion of significant SNPs (P < 5 × 10^−8^) to non-significant SNPs (P > 5 × 10^−8^) within cluster-enriched genes against all genes detected in the scRNA-seq dataset.

Cell-cell interactions were predicted using CellChat (Jin, Plikus et al. 2025) which forecasts potential interactions, employs network and pattern recognition by analyzing predefined ligand-receptor pairs.

### Statistical analyses

All statistical analyses were carried out in R. Unpaired one- or two-sample Student’s t-test were used for quantification of bone formation, ALP activity, and Seahorse measurements. Statistics for sequencing-based methods were derived from Seurat, DEseq2, and limma. Replicates are indicated in the individual figures.

## Acknowledgements

Sequencing was carried out at the Villum Center for Bioanalytical Sciences, Functional Genomics & Metabolism Research Unit, University of Southern Denmark. We thank Ronni Nielsen for sequencing assistance. This work was supported by a grant from the Lundbeck Foundation (R335-2019-2195) and Independent Research Fund Denmark (0134-00292B) to A.R., a grant from the Novo Nordisk Foundation (NNF19OC0057449) to M.K., and a PhD scholarship from the Faculty of Health Science University of Southern Denmark to A.C.

## Author Contribution Statement (CRediT)

**Atenisa Caci:** Formal analysis, Investigation, Data curation, Visualization, Writing – original draft. **Nicholas Ditzel**: Data curation. **Sebastian Zanner**: Data curation. **Moustapha Kassem:** Funding acquisition, Supervision, Writing - review & editing **Alexander Rauch:** Conceptualization, Funding acquisition, Supervision, Writing - review & editing

## Ethics Statement

Ethical approval for animal experiments was given by the Animal Experimentation Council of the Ministry of Food, Agriculture and Fishing Denmark (License 2022-15-0201-01225).

## Data Availability Statement

Raw sequencing data of hBMSC-TERT4 cells reported in this study have been deposited under the NCBI Gene Expression Omnibus: GSEXXX. Scripts for data processing and visualization to recapitulate the analyses of sequencing-based data are available at the Open Science Framework: https://osf.io/wxpgn and at GitHub: https://github.com/atenisacaci/hBMSC-TERT4-clones.

## References

Abdallah, B. M., N. Ditzel and M. Kassem (2008). “Assessment of bone formation capacity using in vivo transplantation assays: procedure and tissue analysis.” Methods Mol Biol 455: 89–100.

Ambrosi, T. H., R. Sinha, H. M. Steininger, M. Y. Hoover, M. P. Murphy, L. S. Koepke, Y. Wang, W. J. Lu, M. Morri, N. F. Neff, I. L. Weissman, M. T. Longaker and C. K. Chan (2021). “Distinct skeletal stem cell types orchestrate long bone skeletogenesis.” Elife 10.

Ambrosi, T. H., S. Taheri, K. Chen, R. Sinha, Y. Wang, E. J. Hunt, L. H. Goodnough, M. P. Murphy, H. M. Steininger, M. Y. Hoover, F. Felix, K. C. Weldon, L. S. Koepke, J. Sokol, D. D. Liu, L. Zhao, S. D. Conley, W. J. Lu, M. Morri, N. F. Neff, N. L. Van Rysselberghe, E. E. Wheeler, Y. Wang, J. K. Leach, A. Saiz, A. Wang, G. P. Yang, S. Goodman, J. A. Bishop, M. J. Gardner, D. C. Wan, I. L. Weissman, M. T. Longaker, D. Sahoo and C. K. F. Chan (2025). “Human skeletal development and regeneration are shaped by functional diversity of stem cells across skeletal sites.” Cell Stem Cell.

Ashburner, M., C. A. Ball, J. A. Blake, D. Botstein, H. Butler, J. M. Cherry, A. P. Davis, K. Dolinski, S. S. Dwight, J. T. Eppig, M. A. Harris, D. P. Hill, L. Issel-Tarver, A. Kasarskis, S. Lewis, J. C. Matese, J. E. Richardson, M. Ringwald, G. M. Rubin and G. Sherlock (2000). “Gene ontology: tool for the unification of biology. The Gene Ontology Consortium.” Nat Genet 25(1): 25–29.

Bandyopadhyay, S., M. P. Duffy, K. J. Ahn, J. H. Sussman, M. Pang, D. Smith, G. Duncan, I. Zhang, J. Huang, Y. Lin, B. Xiong, T. Imtiaz, C. H. Chen, A. Thadi, C. Chen, J. Xu, M. Reichart, Z. Martinez, C. Diorio, C. Chen, V. Pillai, O. Snaith, D. Oldridge, S. Bhattacharyya, I. Maillard, M. Carroll, C. Nelson, L. Qin and K. Tan (2024). “Mapping the cellular biogeography of human bone marrow niches using single-cell transcriptomics and proteomic imaging.” Cell 187(12): 3120–3140 e3129.

Baryawno, N., D. Przybylski, M. S. Kowalczyk, Y. Kfoury, N. Severe, K. Gustafsson, K. D. Kokkaliaris, F. Mercier, M. Tabaka, M. Hofree, D. Dionne, A. Papazian, D. Lee, O. Ashenberg, A. Subramanian, E. D. Vaishnav, O. Rozenblatt-Rosen, A. Regev and D. T. Scadden (2019). “A Cellular Taxonomy of the Bone Marrow Stroma in Homeostasis and Leukemia.” Cell 177(7): 1915–1932 e1916.

Buenrostro, J. D., M. R. Corces, C. A. Lareau, B. Wu, A. N. Schep, M. J. Aryee, R. Majeti, H. Y. Chang and W. J. Greenleaf (2018). “Integrated Single-Cell Analysis Maps the Continuous Regulatory Landscape of Human Hematopoietic Differentiation.” Cell 173(6): 1535–1548 e1516.

Butler, A., P. Hoffman, P. Smibert, E. Papalexi and R. Satija (2018). “Integrating single-cell transcriptomic data across different conditions, technologies, and species.” Nat Biotechnol 36(5): 411–420.

Chai, R. C., M. Lundberg, B. Freudenthal, J. T. Smith, A. P. Boughton, Y. Zhang, K. A. Flynn, M. Frysz, A. P. Corr, W. H. Khoo, D. Komla-Ebri, M. R. G. Dack, S. E. Guilfoyle, J. G. Logan, N. C. Butterfield, V. D. Leitch, A. S. Pollard, R. E. Makitie, N. Bradford, L. Ramos-Mucci, A. Vilas-Zornoza, Y. Chen, R. K. H. Yip, J. Er, S. Z. Tan, M. M. McDonald, S. E. Youlten, C. M. Sergio, A. Castro-Martinez, S. G. Young, E. Skorokhodova, D. M. Evans, J. E. Powell, C. A. de Leeuw, A. D. Ewing, J. A. Eisman, R. D. Blank, T. G. Phan G. International Federation of Musculoskeletal Research Societies Big Data Working, A. K. Lagendijk, E. D. Hawkins, H. L. Oh, R. E. McIntyre, E. M. Hessel, J. P. Taylor-King, P. A. Baldock, E. L. Duncan, G. R. Williams, J. H. D. Bassett, P. I. Croucher and J. P. Kemp (2026). “Multiscale analysis and functional validation of the cellular and genetic determinants of skeletal disease.” Nat Genet 58(7): 1530–1545.

Chen, C. T., Y. R. Shih, T. K. Kuo, O. K. Lee and Y. H. Wei (2008). “Coordinated changes of mitochondrial biogenesis and antioxidant enzymes during osteogenic differentiation of human mesenchymal stem cells.” Stem Cells 26(4): 960–968.

Collins, J. M., A. Lang, C. Parisi, Y. Moharrer, M. P. Nijsure, J. H. Thomas Kim, S. Ahmed, G. L. Szeto, L. Qin, R. Gottardi, N. A. Dyment, N. C. Nowlan and J. D. Boerckel (2024). “YAP and TAZ couple osteoblast precursor mobilization to angiogenesis and mechanoregulation in murine bone development.” Dev Cell 59(2): 211–227 e215.

Davis, S. and P. S. Meltzer (2007). “GEOquery: a bridge between the Gene Expression Omnibus (GEO) and BioConductor.” Bioinformatics 23(14): 1846–1847.

Debnath, S., A. R. Yallowitz, J. McCormick, S. Lalani, T. Zhang, R. Xu, N. Li, Y. Liu, Y. S. Yang, M. Eiseman, J. H. Shim, M. Hameed, J. H. Healey, M. P. Bostrom, D. A. Landau and M. B. Greenblatt (2018). “Discovery of a periosteal stem cell mediating intramembranous bone formation.” Nature 562(7725): 133–139.

Deveza, L., L. Ortinau, K. Lei and D. Park (2018). “Comparative analysis of gene expression identifies distinct molecular signatures of bone marrow- and periosteal-skeletal stem/progenitor cells.” PLoS One 13(1): e0190909.

Dominici, M., K. Le Blanc, I. Mueller, I. Slaper-Cortenbach, F. Marini, D. Krause, R. Deans, A. Keating, D. Prockop and E. Horwitz (2006). “Minimal criteria for defining multipotent mesenchymal stromal cells. The International Society for Cellular Therapy position statement.” Cytotherapy 8(4): 315–317.

Dupont, S., L. Morsut, M. Aragona, E. Enzo, S. Giulitti, M. Cordenonsi, F. Zanconato, J. Le Digabel, M. Forcato, S. Bicciato, N. Elvassore and S. Piccolo (2011). “Role of YAP/TAZ in mechanotransduction.” Nature 474(7350): 179–183.

Gautier, L., L. Cope, B. M. Bolstad and R. A. Irizarry (2004). “affy--analysis of Affymetrix GeneChip data at the probe level.” Bioinformatics 20(3): 307–315.

Guo, L., T. Cai, K. Chen, R. Wang, J. Wang, C. Cui, J. Yuan, K. Zhang, Z. Liu, Y. Deng, G. Xiao and C. Wu (2018). “Kindlin-2 regulates mesenchymal stem cell differentiation through control of YAP1/TAZ.” J Cell Biol 217(4): 1431–1451.

Hong, J. H., E. S. Hwang, M. T. McManus, A. Amsterdam, Y. Tian, R. Kalmukova, E. Mueller, T. Benjamin, B. M. Spiegelman, P. A. Sharp, N. Hopkins and M. B. Yaffe (2005). “TAZ, a transcriptional modulator of mesenchymal stem cell differentiation.” Science 309(5737): 1074–1078.

Jamil, A. J. M., K. Madsen, S. Daamouch, B. Villadsen, C. Ma, S. Jeromdesella, M. O. Norgard, E. Grupe, M. B. Jakobsson, Y. Wang, E. Takematsu, L. Zhao, S. B. Goodman, L. H. Goodnough, C. K. F. Chan, T. L. Andersen, M. Kassem and A. Rauch (2026). “An ITGA11 expressing subpopulation as predictor for the donor-specific osteogenic capacity of stromal cells.” Bone Res 14(1).

Janicki, P., S. Boeuf, E. Steck, M. Egermann, P. Kasten and W. Richter (2011). “Prediction of in vivo bone forming potency of bone marrow-derived human mesenchymal stem cells.” Eur Cell Mater 21: 488–507.

Jeffery, E. C., T. L. A. Mann, J. A. Pool, Z. Zhao and S. J. Morrison (2022). “Bone marrow and periosteal skeletal stem/progenitor cells make distinct contributions to bone maintenance and repair.” Cell Stem Cell 29(11): 1547–1561 e1546.

Jemtland, R., M. Holden, S. Reppe, O. K. Olstad, F. P. Reinholt, V. T. Gautvik, H. Refvem, A. Frigessi, B. Houston and K. M. Gautvik (2011). “Molecular disease map of bone characterizing the postmenopausal osteoporosis phenotype.” J Bone Miner Res 26(8): 1793–1801.

Jin, S., M. V. Plikus and Q. Nie (2025). “CellChat for systematic analysis of cell-cell communication from single-cell transcriptomics.” Nat Protoc 20(1): 180–219.

Kastan, N., K. Gnedeva, T. Alisch, A. A. Petelski, D. J. Huggins, J. Chiaravalli, A. Aharanov, A. Shakked, E. Tzahor, A. Nagiel, N. Segil and A. J. Hudspeth (2021). “Small-molecule inhibition of Lats kinases may promote Yap-dependent proliferation in postmitotic mammalian tissues.” Nat Commun 12(1): 3100.

Kegelman, C. D., M. P. Nijsure, Y. Moharrer, H. B. Pearson, J. H. Dawahare, K. M. Jordan, L. Qin and J. D. Boerckel (2021). “YAP and TAZ Promote Periosteal Osteoblast Precursor Expansion and Differentiation for Fracture Repair.” J Bone Miner Res 36(1): 143–157.

Kowal, J. M., S. Moller, D. Ali, F. Figeac, T. Barington, H. Schmal and M. Kassem (2021). “Identification of a clinical signature predictive of differentiation fate of human bone marrow stromal cells.” Stem Cell Res Ther 12(1): 265.

Kowal, J. M., H. Schmal, U. Halekoh, J. B. Hjelmborg and M. Kassem (2020). “Single-cell high-content imaging parameters predict functional phenotype of cultured human bone marrow stromal stem cells.” Stem Cells Transl Med 9(2): 189–202.

Larsen, K. H., C. M. Frederiksen, J. S. Burns, B. M. Abdallah and M. Kassem (2010). “Identifying a molecular phenotype for bone marrow stromal cells with in vivo bone-forming capacity.” J Bone Miner Res 25(4): 796–808.

Lawrence, M., W. Huber, H. Pages, P. Aboyoun, M. Carlson, R. Gentleman, M. T. Morgan and V. J. Carey (2013). “Software for computing and annotating genomic ranges.” PLoS Comput Biol 9(8): e1003118.

Li, Q., R. Xu, K. Lei and Q. Yuan (2022). “Insights into skeletal stem cells.” Bone Res 10(1): 61.

Liao, X., J. Koehnken Sawall, R. Seeley, F. Song, X. Ji, X. Liu, C. Song and F. Long (2025). “An adipoosteoprogenitor population in the endosteal niche contributes to bone and fat formation in adult mouse bone marrow.” Proc Natl Acad Sci U S A 122(47): e2502436122.

Liu, H., P. Li, S. Zhang, J. Xiang, R. Yang, J. Liu, M. Shafiquzzaman, S. Biswas, Z. Wei, Z. Zhang, X. Zhou, F. Yin, Y. Xie, S. P. Goff, L. Chen and B. Li (2022). “Prrx1 marks stem cells for bone, white adipose tissue and dermis in adult mice.” Nat Genet 54(12): 1946–1958.

Liu, X., Y. Ye, Z. Li, L. Liao and Q. Wei (2026). “Mechanical rejuvenation of senescent stem cells and aged bone via chromatin remodeling.” Nat Commun 17(1): 1684.

Maes, C., T. Kobayashi and H. M. Kronenberg (2007). “A novel transgenic mouse model to study the osteoblast lineage in vivo.” Ann N Y Acad Sci 1116: 149–164.

Matsushita, Y., M. Nagata, K. M. Kozloff, J. D. Welch, K. Mizuhashi, N. Tokavanich, S. A. Hallett, D. C. Link, T. Nagasawa, W. Ono and N. Ono (2020). “A Wnt-mediated transformation of the bone marrow stromal cell identity orchestrates skeletal regeneration.” Nat Commun 11(1): 332.

Matthews, B. G., N. K. Y. Wee, V. N. Widjaja, J. S. Price, I. Kalajzic and S. H. Windahl (2020). “alphaSMA Osteoprogenitor Cells Contribute to the Increase in Osteoblast Numbers in Response to Mechanical Loading.” Calcif Tissue Int 106(2): 208–217.

Melis, S., D. Trompet, A. S. Chagin and C. Maes (2025). “Skeletal stem and progenitor cells in bone physiology, ageing and disease.” Nat Rev Endocrinol 21(3): 135–153.

Mendez-Ferrer, S., D. Bonnet, D. P. Steensma, R. P. Hasserjian, I. M. Ghobrial, J. G. Gribben, M. Andreeff and D. S. Krause (2020). “Bone marrow niches in haematological malignancies.” Nat Rev Cancer 20(5): 285–298.

Mendez-Ferrer, S., T. V. Michurina, F. Ferraro, A. R. Mazloom, B. D. Macarthur, S. A. Lira, D. T. Scadden, A. Ma’ayan, G. N. Enikolopov and P. S. Frenette (2010). “Mesenchymal and haematopoietic stem cells form a unique bone marrow niche.” Nature 466(7308): 829–834.

Meuleman, W., A. Muratov, E. Rynes, J. Halow, K. Lee, D. Bates, M. Diegel, D. Dunn, F. Neri, A. Teodosiadis, A. Reynolds, E. Haugen, J. Nelson, A. Johnson, M. Frerker, M. Buckley, R. Sandstrom, J. Vierstra, R. Kaul and J. Stamatoyannopoulos (2020). “Index and biological spectrum of human DNase I hypersensitive sites.” Nature 584(7820): 244–251.

Mizuhashi, K., W. Ono, Y. Matsushita, N. Sakagami, A. Takahashi, T. L. Saunders, T. Nagasawa, H. M. Kronenberg and N. Ono (2018). “Resting zone of the growth plate houses a unique class of skeletal stem cells.” Nature 563(7730): 254–258.

Mo, C., J. Guo, J. Qin, X. Zhang, Y. Sun, H. Wei, D. Cao, Y. Zhang, C. Zhao, Y. Xiong, Y. Zhang, Y. Sun, L. Shen and R. Yue (2022). “Single-cell transcriptomics of LepR-positive skeletal cells reveals heterogeneous stress-dependent stem and progenitor pools.” EMBO J 41(4): e108415.

Moller, A. F. and J. G. S. Madsen (2023). “JOINTLY: interpretable joint clustering of single-cell transcriptomes.” Nat Commun 14(1): 8473.

Morris, J. A., J. P. Kemp, S. E. Youlten, L. Laurent, J. G. Logan, R. C. Chai, N. A. Vulpescu, V. Forgetta, A. Kleinman, S. T. Mohanty, C. M. Sergio, J. Quinn, L. Nguyen-Yamamoto, A. L. Luco, J. Vijay, M. M. Simon, A. Pramatarova, C. Medina-Gomez, K. Trajanoska, E. J. Ghirardello, N. C. Butterfield, K. F. Curry, V. D. Leitch, P. C. Sparkes, A. T. Adoum, N. S. Mannan, D. S. K. Komla-Ebri, A. S. Pollard, H. F. Dewhurst, T. A. D. Hassall, M. G. Beltejar, T. andMe Research, D. J. Adams, S. M. Vaillancourt, S. Kaptoge, P. Baldock, C. Cooper, J. Reeve, E. E. Ntzani, E. Evangelou, C. Ohlsson, D. Karasik, F. Rivadeneira, D. P. Kiel, J. H. Tobias, C. L. Gregson, N. C. Harvey, E. Grundberg, D. Goltzman, D. J. Adams, C. J. Lelliott, D. A. Hinds, C. L. Ackert-Bicknell, Y. H. Hsu, M. T. Maurano, P. I. Croucher, G. R. Williams, J. H. D. Bassett, D. M. Evans and J. B. Richards (2019). “An atlas of genetic influences on osteoporosis in humans and mice.” Nat Genet 51(2): 258–266.

Nguyen, H. C. T., B. Baik, S. Yoon, T. Park and D. Nam (2023). “Benchmarking integration of single-cell differential expression.” Nat Commun 14(1): 1570.

Nijsure, M. P., B. Tobin, D. L. Jones, A. Lang, G. Hallstrom, M. Baitner, G. I. Tanner, G. Kotsaris, Y. Moharrer, C. J. Panebianco, E. G. Seidl, N. A. Dyment, G. L. Szeto, L. Wood and J. D. Boerckel (2025). “YAP regulates transcriptional programs for layer-specific periosteal expansion during fracture repair.” Sci Adv 11(32): eadw0126.

Nouri, K., T. Azad, M. Ling, H. J. Janse van Rensburg, A. Pipchuk, H. Shen, Y. Hao, J. Zhang and X. Yang (2019). “Identification of Celastrol as a Novel YAP-TEAD Inhibitor for Cancer Therapy by High Throughput Screening with Ultrasensitive YAP/TAZ-TEAD Biosensors.” Cancers (Basel) 11(10).

Omatsu, Y., T. Sugiyama, H. Kohara, G. Kondoh, N. Fujii, K. Kohno and T. Nagasawa (2010). “The essential functions of adipo-osteogenic progenitors as the hematopoietic stem and progenitor cell niche.” Immunity 33(3): 387–399.

Ono, N., W. Ono, T. Nagasawa and H. M. Kronenberg (2014). “A subset of chondrogenic cells provides early mesenchymal progenitors in growing bones.” Nat Cell Biol 16(12): 1157–1167.

Panciera, T., L. Azzolin, M. Cordenonsi and S. Piccolo (2017). “Mechanobiology of YAP and TAZ in physiology and disease.” Nat Rev Mol Cell Biol 18(12): 758–770.

Parekh, S., C. Ziegenhain, B. Vieth, W. Enard and I. Hellmann (2018). “zUMIs - A fast and flexible pipeline to process RNA sequencing data with UMIs.” Gigascience 7(6).

Pico, A. R., T. Kelder, M. P. van Iersel, K. Hanspers, B. R. Conklin and C. Evelo (2008). “WikiPathways: pathway editing for the people.” PLoS Biol 6(7): e184.

Ragueneau, E., C. Gong, P. Sinquin, C. Sevilla, D. Beavers, A. Grentner, J. Griss, G. F. J. Hogue, N. T. Li, L. Matthews, B. May, M. Milacic, H. Mohammadi, R. Petryszak, K. Rothfels, V. Shamovsky, R. Stephan, K. Tiwari, J. Weiser, A. Wright, M. Gillespie, G. Wu, L. Stein, H. Hermjakob and P. D’Eustachio (2026). “The Reactome Knowledgebase 2026.” Nucleic Acids Res 54(D1): D673–D681.

Rauch, A., A. K. Haakonsson, J. G. S. Madsen, M. Larsen, I. Forss, M. R. Madsen, E. L. Van Hauwaert, C. Wiwie, N. Z. Jespersen, M. Tencerova, R. Nielsen, B. D. Larsen, R. Rottger, J. Baumbach, C. Scheele, M. Kassem and S. Mandrup (2019). “Osteogenesis depends on commissioning of a network of stem cell transcription factors that act as repressors of adipogenesis.” Nat Genet 51(4): 716–727.

Rauch, A. and S. Mandrup (2021). “Transcriptional networks controlling stromal cell differentiation.” Nat Rev Mol Cell Biol 22(7): 465–482.

Ritchie, M. E., B. Phipson, D. Wu, Y. Hu, C. W. Law, W. Shi and G. K. Smyth (2015). “limma powers differential expression analyses for RNA-sequencing and microarray studies.” Nucleic Acids Res 43(7): e47.

Rydenfelt, M., B. Klinger, M. Klünemann and N. Blüthgen (2020). “SPEED2: inferring upstream pathway activity from differential gene expression.” Nucleic Acids Research 48(W1): W307–W312.

Shen, B., A. Tasdogan, J. M. Ubellacker, J. Zhang, E. D. Nosyreva, L. Du, M. M. Murphy, S. Hu, Y. Yi, N. Kara, X. Liu, S. Guela, Y. Jia, V. Ramesh, C. Embree, E. C. Mitchell, Y. C. Zhao, L. A. Ju, Z. Hu, G. M. Crane, Z. Zhao, R. Syeda and S. J. Morrison (2021). “A mechanosensitive peri-arteriolar niche for osteogenesis and lymphopoiesis.” Nature 591(7850): 438–444.

Shi, Y., G. He, W. C. Lee, J. A. McKenzie, M. J. Silva and F. Long (2017). “Gli1 identifies osteogenic progenitors for bone formation and fracture repair.” Nat Commun 8(1): 2043.

Simonsen, J. L., C. Rosada, N. Serakinci, J. Justesen, K. Stenderup, S. I. Rattan, T. G. Jensen and M. Kassem (2002). “Telomerase expression extends the proliferative life-span and maintains the osteogenic potential of human bone marrow stromal cells.” Nat Biotechnol 20(6): 592–596.

Simonsen, J. L., C. Rosada, N. Serakinci, J. Justesen, K. Stenderup, S. I. S. Rattan, T. G. Jensen and M. Kassem (2002). “Telomerase expression extends the proliferative life-span and maintains the osteogenic potential of human bone marrow stromal cells.” Nature Biotechnology 20(6): 592–596.

Stergachis, A. B., S. Neph, A. Reynolds, R. Humbert, B. Miller, S. L. Paige, B. Vernot, J. B. Cheng, R. E. Thurman, R. Sandstrom, E. Haugen, S. Heimfeld, C. E. Murry, J. M. Akey and J. A. Stamatoyannopoulos (2013). “Developmental fate and cellular maturity encoded in human regulatory DNA landscapes.” Cell 154(4): 888–903.

Tikhonova, A. N., I. Dolgalev, H. Hu, K. K. Sivaraj, E. Hoxha, A. Cuesta-Dominguez, S. Pinho, I. Akhmetzyanova, J. Gao, M. Witkowski, M. Guillamot, M. C. Gutkin, Y. Zhang, C. Marier, C. Diefenbach, S. Kousteni, A. Heguy, H. Zhong, D. R. Fooksman, J. M. Butler, A. Economides, P. S. Frenette, R. H. Adams, R. Satija, A. Tsirigos and I. Aifantis (2019). “The bone marrow microenvironment at single-cell resolution.” Nature 569(7755): 222–228.

Trompet, D., S. Melis, A. S. Chagin and C. Maes (2024). “Skeletal stem and progenitor cells in bone development and repair.” J Bone Miner Res 39(6): 633–654.

Walzl, A., C. Unger, N. Kramer, D. Unterleuthner, M. Scherzer, M. Hengstschlager, D. Schwanzer-Pfeiffer and H. Dolznig (2014). “The Resazurin Reduction Assay Can Distinguish Cytotoxic from Cytostatic Compounds in Spheroid Screening Assays.” Journal of Biomolecular Screening 19(7): 1047–1059.

Wang, Z., P. Liu, X. Zhou, T. Wang, X. Feng, Y. P. Sun, Y. Xiong, H. X. Yuan and K. L. Guan (2017). “Endothelin Promotes Colorectal Tumorigenesis by Activating YAP/TAZ.” Cancer Res 77(9): 2413–2423.

Weinreb, C., A. Rodriguez-Fraticelli, F. D. Camargo and A. M. Klein (2020). “Lineage tracing on transcriptional landscapes links state to fate during differentiation.” Science 367(6479).

Weldon, K. C., M. T. Longaker and T. H. Ambrosi (2025). “Harnessing the diversity and potential of endogenous skeletal stem cells for musculoskeletal tissue regeneration.” Stem Cells 43(3).

Worthley, D. L., M. Churchill, J. T. Compton, Y. Tailor, M. Rao, Y. Si, D. Levin, M. G. Schwartz, A. Uygur, Y. Hayakawa, S. Gross, B. W. Renz, W. Setlik, A. N. Martinez, X. Chen, S. Nizami, H. G. Lee, H. P. Kang, J. M. Caldwell, S. Asfaha, C. B. Westphalen, T. Graham, G. Jin, K. Nagar, H. Wang, M. A. Kheirbek, A. Kolhe, J. Carpenter, M. Glaire, A. Nair, S. Renders, N. Manieri, S. Muthupalani, J. G. Fox, M. Reichert, A. S. Giraud, R. F. Schwabe, J. P. Pradere, K. Walton, A. Prakash, D. Gumucio, A. K. Rustgi, T. S. Stappenbeck, R. A. Friedman, M. D. Gershon, P. Sims, T. Grikscheit, F. Y. Lee, G. Karsenty, S. Mukherjee and T. C. Wang (2015). “Gremlin 1 identifies a skeletal stem cell with bone, cartilage, and reticular stromal potential.” Cell 160(1-2): 269–284.

Wu, S., H. Matsumoto, K. Kondo, A. Kuroda, J. Morita, T. Moriishi, S. A. Hallett, Y. Matsuo, A. Noguchi, M. Ono, M. Abe, K. Ito, S. Ohba, N. Ono and Y. Matsushita (2026). “Injury-driven stromal exhaustion disrupts intrinsic regenerative capability.” Nat Commun 17(1).

Xiong, J., M. Almeida and C. A. O’Brien (2018). “The YAP/TAZ transcriptional co-activators have opposing effects at different stages of osteoblast differentiation.” Bone 112: 1–9.

Yang, R., D. Cao, J. Suo, L. Zhang, C. Mo, M. Wang, N. Niu, R. Yue and W. Zou (2023). “Premature aging of skeletal stem/progenitor cells rather than osteoblasts causes bone loss with decreased mechanosensation.” Bone Res 11(1): 35.

Young, M. D., M. J. Wakefield, G. K. Smyth and A. Oshlack (2010). “Gene ontology analysis for RNA-seq: accounting for selection bias.” Genome Biol 11(2): R14.

Yu, F. X., B. Zhao and K. L. Guan (2015). “Hippo Pathway in Organ Size Control, Tissue Homeostasis, and Cancer.” Cell 163(4): 811–828.

Zhong, L., L. Yao, R. J. Tower, Y. Wei, Z. Miao, J. Park, R. Shrestha, L. Wang, W. Yu, N. Holdreith, X. Huang, Y. Zhang, W. Tong, Y. Gong, J. Ahn, K. Susztak, N. Dyment, M. Li, F. Long, C. Chen, P. Seale and L. Qin (2020). “Single cell transcriptomics identifies a unique adipose lineage cell population that regulates bone marrow environment.” Elife 9.

Zhou, B. O., R. Yue, M. M. Murphy, J. G. Peyer and S. J. Morrison (2014). “Leptin-receptor-expressing mesenchymal stromal cells represent the main source of bone formed by adult bone marrow.” Cell Stem Cell 15(2): 154–168.

Zhu, Q., L. Ding and R. Yue (2022). “Skeletal stem cells: a game changer of skeletal biology and regenerative medicine?” Life Med 1(3): 294–306.

Zhu, Y., Y. Wu, J. Cheng, Q. Wang, Z. Li, Y. Wang, D. Wang, H. Wang, W. Zhang, J. Ye, H. Jiang and L. Wang (2018). “Pharmacological activation of TAZ enhances osteogenic differentiation and bone formation of adipose-derived stem cells.” Stem Cell Res Ther 9(1): 53.

